# Layer-dependent loss and enhancement of geniculostriate and retinotectal pathways in adult human amblyopia

**DOI:** 10.1101/2020.12.29.424669

**Authors:** Wen Wen, Yue Wang, Sheng He, Hong Liu, Chen Zhao, Peng Zhang

**Affiliations:** Department of Ophthalmology & Visual Science, Eye & ENT Hospital, Shanghai Medical College, Fudan University, Shanghai, China; State Key Laboratory of Brain and Cognitive Science, Institute of Biophysics, Chinese Academy of Sciences, Beijing, 100101, China; University of Chinese Academy of Sciences, Beijing 100049, China; Institute of Artificial Intelligence, Hefei Comprehensive National Science Center, Hefei 230026, China; Department of Psychology, University of Minnesota, Minneapolis, Minnesota, USA; State Key Laboratory of Medical Neurobiology, Institutes of Brain Science, Fudan University, Shanghai, China; Key Laboratory of Myopia, Ministry of Health, Fudan University, Shanghai, China; Shanghai Key Laboratory of Visual Impairment and Restoration, Fudan University, Shanghai, China; Department of Ophthalmology, Shanghai Children’s Medical Center, Shanghai Jiao Tong University School of Medicine, Shanghai, China

**Keywords:** Layers, fMRI, LGN, Magnocellular, Parvocellular, SC, Pulvinar

## Abstract

Abnormal visual experience during critical period leads to reorganization of neuroarchitectures in primate visual cortex. However, developmental plasticity of human subcortical visual pathways remains elusive. Using high-resolution fMRI and pathway-selective visual stimuli, we investigated layer-dependent response properties and connectivity of subcortical visual pathways of adult human amblyopia. Stimuli presented to the amblyopic eye showed selective response loss in the parvocellular layers of the lateral geniculate nucleus, and also reduced the connectivity to V1. Amblyopic eye’s response to isoluminant chromatic stimulus was significantly reduced in the superficial layers of the superior colliculus, while the fellow eye’s response robustly increased in the deeper layers associated with increased cortical feedbacks. Therefore, amblyopia led to selective reduction of parvocellular feedforward signals in the geniculostriate pathway, whereas loss and enhancement of parvocellular feedback signals in the retinotectal pathway. These findings shed light for future development of new tools for treating amblyopia and tracking the prognosis.

**Highlights:**

- Amblyopia impairs feedforward processing in the P layers of the LGN
- Layer-dependent loss and enhancement of cortical feedback signals in the SC
- Pathway-specific abnormalities explain amblyopic deficits in visual acuity and attention

**Significance statement:** How abnormal visual experiences during critical period shape the function and wire the neural circuits of human subcortex remains largely unknown. With high-resolution fMRI and visual stimuli to preferentially activate layer-dependent response in human subcortical pathways, the current study clearly demonstrates that amblyopia shifts the homeostatic interocular balance of human subcortex in a pathway-specific manner. Amblyopia led to selective loss of parvocellular feedforward signals in the geniculostriate pathway, whereas deficit and enhancement of parvocellular feedback signals in the retinotectal pathway. These pathwayspecific functional abnormalities provide the neural basis for amblyopic deficits in visual acuity, control of eye movement and attention. It sheds light for future development of new tools for treating amblyopia and tracking the prognosis.

## Introduction

Abnormal visual experience in the critical period, commonly due to a turned eye (strabismus) or unequal refractive errors (anisometropia), causes amblyopia or lazy eye characterized by impaired visual abilities even with corrected optics. Amblyopia serves as a model to understand how abnormal visual experience during critical period reshapes the wiring and functions of the human visual system. Retinal structures and functions of the amblyopic eye have been found to be intact (Hess & Baker Jr., 1984; Levi, 2006). Animal models showed shrinkage of ocular dominance columns in the primary visual cortex and cell atrophy in the deprived eye’s layers of the lateral geniculate nucleus (LGN) (Hubel et al., 1977; Movshon et al., 1987; Wiesel & Hubel, 1963). In the human brain, neuroimaging studies found reduced response to the amblyopic eye in the visual cortex (Anderson & Swettenham, 2006; Joly & Franko, 2014). Due to technical limitations to access layer-specific activity of human subcortical nuclei, much less is known about the influence of amblyopia on the neural response and circuits of human subcortical visual pathways.

In primate, retinal information reaches visual cortex mainly through the geniculostriate pathway. The lateral geniculate nucleus (LGN) is composed of six main layers of neurons with distinct cell types and functions. Parvocellular (P) cells in the four dorsal layers have smaller cell body, sustained discharge pattern, smaller receptive field, low sensitivity to luminance contrast, and center-surround color opponency; magnocellular (M) cells in the two ventral layers have larger cell body, transient neural response, larger receptive field, and high sensitivity to luminance contrast and motion (Derrington et al., 1984; Derrington & Lennie, 1984; Hubel & Livingstone, 1990). Thus, the M and P stream-specific visual processing are clearly segregated in the LGN, which serves as a perfect brain area to study the M and/or P selective deficits in amblyopia. However, lacking a non-invasive tool to directly access layer-specific response of the human LGN, existing evidence from human psychophysics and neuroimaging of visual cortex is controversial (Levi and Harwerth 1977, Barnes, Hess et al. 2001, Zele, Pokorny et al. 2007, Hess, Li et al. 2009). The disagreement from the previous studies are likely due to large differences in visual stimuli, which greatly differed in selectivity to the M and P visual streams. Furthermore, most of the neuroimaging studies measured responses from the visual cortex, where the two pathways’ signals merge thus cannot be clearly segregated.

Retinotectal pathway is an alternative and more primitive visual pathway in primate. About 10% of retinal ganglion cells project to the superficial layers of the superior colliculus (SC) (Perry & Cowey, 1984), which sends projections to the Koniocellular layers of the LGN and the inferior or visual pulvinar (May, 2006). The superficial layers of the SC (superficial SC, or SCs) also receives direct cortical input from the early visual cortex (V1-V3) and area MT (Fries, 1984; Lock et al., 2003), mainly processes visual sensory information. Deeper layers of the SC (deep SC, or SCd) receive multisensory input from the brain stem (Jay & Sparks, 1987; Wiberg et al., 1987), and afference from the frontoparietal cortex, basal ganglia and extrastriate visual cortex (Beckstead & Frankfurter, 1982; Fries, 1984). The deep SC mainly contributes to attention and premotor control of eye and head movements. The inferior (visual) pulvinar that receives input from the SC sends projections to cortical targets in occipital, parietal and temporal lobes (May, 2006). It plays important roles in visual perception and attention, and regulates information transmission between cortical areas (Purushothaman et al., 2012; Saalmann et al., 2012; Zhou et al., 2016). Amblyopic eyes, especially those with strabismus, are known to exhibit unsteady fixational eye movement (Chung et al., 2015), and several lines of evidence also point to attentional deficits (Verghese et al., 2019). Since the tectal and pulvinar pathways are closely related to the control of eye movement and attention, amblyopic brain may have functional abnormality in these subcortical pathways. Until now, the influence of amblyopia to the tectal and pulvinar pathways remains largely unexplored, thus no direct evidence supports this hypothesis.

Investigating the influence of amblyopia on subcortical responses could reveal how abnormal visual experience during critical period shape the function and wire the circuits of subcortical pathways. Knowing the precise neural deficits is important for developing new tools for better treatment and prognosis of the disease. Recent advances in high resolution fMRI techniques and experimental approaches enable us to resolve layer-specific functional responses of the human LGN and SC (Qian et al., 2020; Yu et al., 2016; Zhang, Wen, et al., 2015; Zhang, Zhou, et al., 2015). In the current study, using high-resolution 3T and 7T BOLD (blood oxygen level dependent) fMRI and carefully designed visual stimuli selectively activating the M and P layers of the LGN, we investigated pathway-specific response and connectivity of the subcortical nuclei of adult amblyopia patients and normal controls.

Our results clearly demonstrated that amblyopia selectively reduced feedforward response in the P layers of the LGN and decreased the effective connectivity from LGN to V1. Amblyopia also led to reduced parvocellular responses in the visual pulvinar, early visual cortex, ventral occipital temporal cortex, and decreased the effective connectivity from pulvinar to V4. Compared to the control eyes, amblyopia eyes’ response to the chromatic P-biased stimulus was significantly reduced in the superficial SC; whereas the response in the deep SC significantly increased to chromatic stimulus presented to the fellow eye, associated with increased parvocellular feedback connectivity from V1. These findings revealed selective loss and enhancement to parvocellular response and connectivity in the subcortical visual pathways of human amblyopia.

## Results

### 1. Amblyopic eye’s response was selectively reduced in the P layers of the LGN

In the 3T fMRI experiment (2mm isotropic voxels), M and P biased visual stimuli (Figure 1A) differed in spatial frequency, temporal frequency, contrast and chromaticity were used to selectively activate the M and P layers of the LGN. The amblyopic eye (AE) and the fellow eye (FE) of unilateral amblyopia patients were tested in separate sessions. The nondominant eye of normal controls (NE) was also tested. We found selective loss of amblyopia eye’s response to the chromatic P stimulus in the P layers of the LGN. In order to reduce the partial volume effect from ventral pulvinar and to further confirm the P layer deficits from the 3T experiment, we performed a 7T fMRI experiment at a higher spatial resolution (1.2mm isotropic voxels). The P cells of the LGN are responsive to both chromatic stimulus and high spatial frequency achromatic stimulus. If there is a general functional loss for the P cells, the P layers’ response to high spatial frequency achromatic stimulus should also be reduced. To test this hypothesis in the 7T experiment, high contrast and high spatial frequency achromatic stimulus (achromatic P stimulus, Figure 2A) was monocularly presented to the amblyopic eye and to the fellow eye of amblyopia patients. The achromatic P stimulus presented to the fellow eye strongly and selectively activated the P layers of the LGN. Compared to the fellow eye, the amblyopic eye’s response was significantly reduced in the P layers of the LGN.

**Figure 1.**
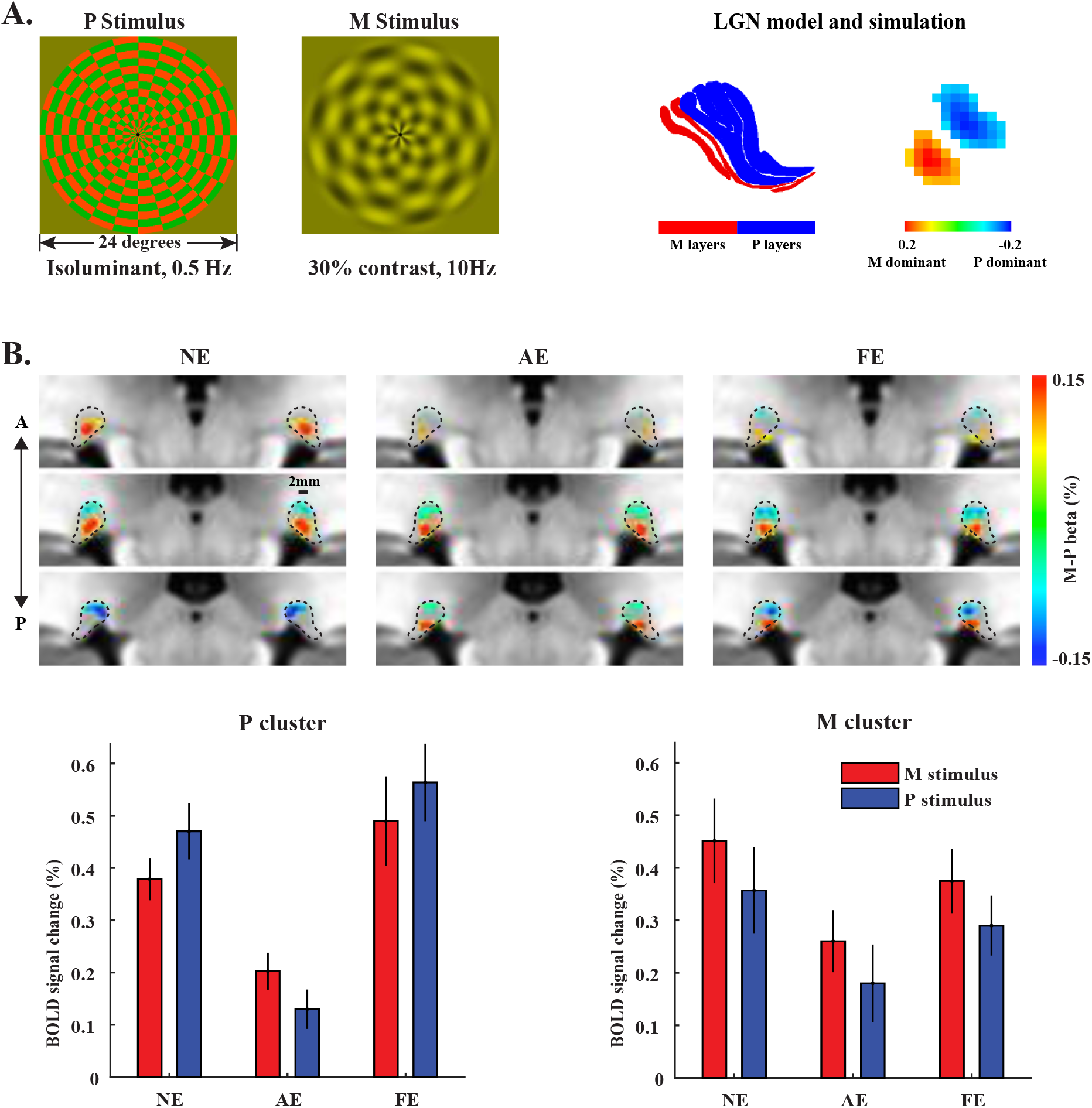
Visual stimuli and results for the 3T fMRI experiment. **A. Left:** M and P biased visual stimuli used in the 3T fMRI experiment. **Right:** The left figure shows the model for the M and P layers of the LGN according to Nissl stained histology of the human LGN (Amunts, Lepage et al. 2013). The right figure shows the simulated fMRI pattern for M-P responses. **B. Upper**: M-P beta maps for the M and P dominant clusters, averaged across all subjects for each group. Beta maps were up-sampled to 0.6mm isotropic resolution for illustration purposes. Maps for the left and right LGNs are mirror symmetric. **Lower:** BOLD responses to the M and P stimuli in the M and P dominant clusters of the LGN. A leave-one-subject-out procedure was used for ROI definition to avoid “double dipping” (see main text for details). Error bars represent standard error of the mean. Abbreviations, NE: normal eye, AE: Amblyopic eye, FE: Fellow eye.

**Figure 2.**
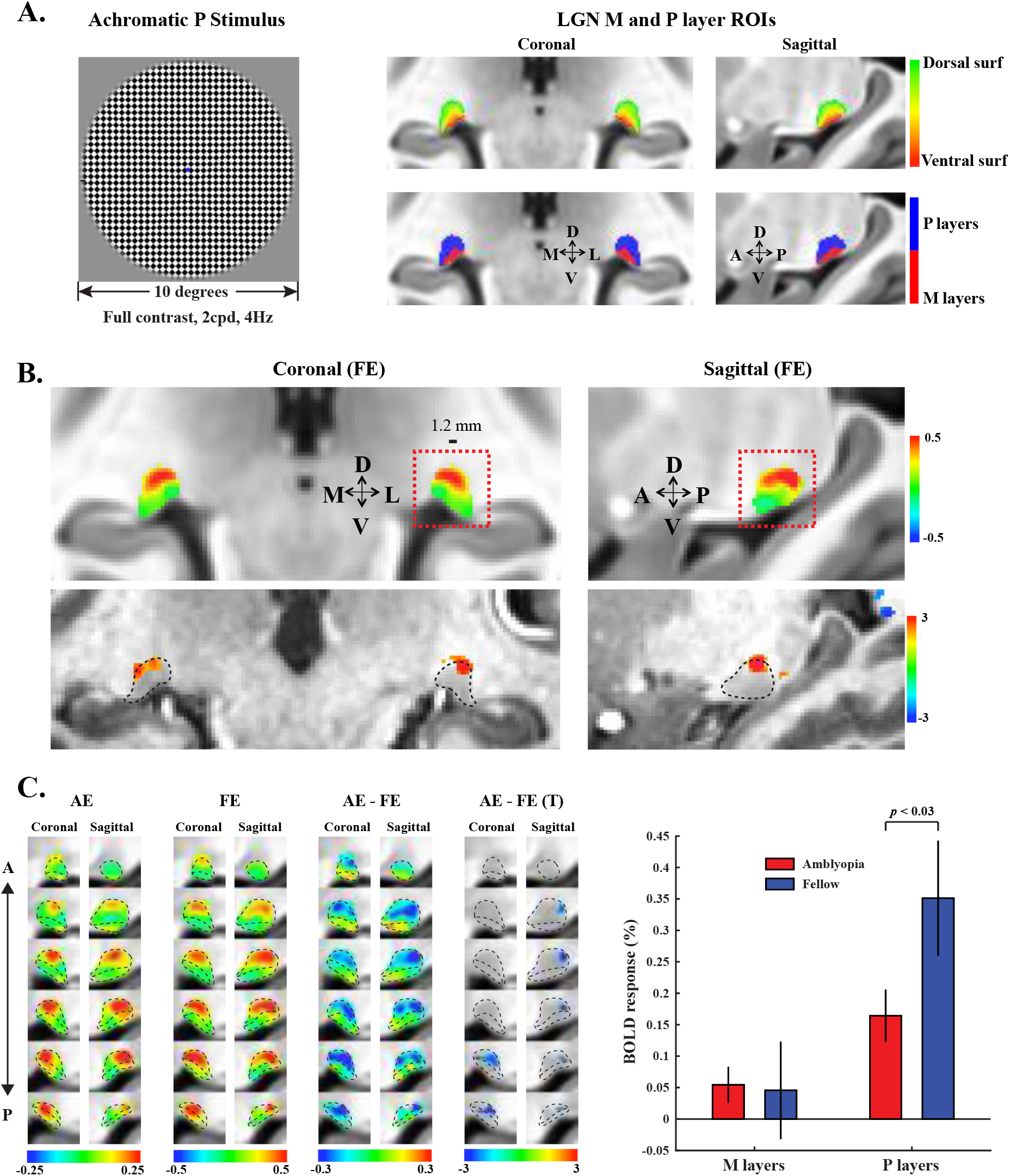
Stimulus and LGN results for the 7T experiment. **A. Left**: Visual stimuli used in the 7T fMRI experiment. **Right**: ROI definitions for the M and P layers of the LGN. The upper panel shows the map for the layer index, calculated as the normalized Euclidean distance to the dorsal and ventral surfaces of the LGN. The lower panels show the ROIs for the M and P layers. **B. Upper:** The high contrast achromatic P stimulus presented to the fellow eye clearly activated the dorsal (P) layers of the LGN (Group averaged beta map). Red boxes indicate the selected FOV for slices showing in C. **Lower:** significant T maps from a representative subject *(p* < 0.05 uncorrected). **C. Left:** The left three columns show the group averaged beta maps for the amblyopic eye (AE), the fellow eye (FE), and the response difference between the AE and NE conditions. The right column shows the T map of significant AE-NE voxels *(p* < 0.05 uncorrected). Maps were up-sampled to 0.6mm isotropic resolution. Black dotted lines indicate the boundary of the M and P layer compartments. **Right:** Amblyopic and fellow eyes’ response in the M and P layers’ ROIs. Error bars represent standard deviation of bootstrapped distribution of the mean.

#### 1.1 3T fMRI showed selective reduction of the amblyopic eye’s response to sustained chromatic stimulus in the P layers of the LGN

As shown by Figure 1A, the chromatic P stimulus was a high spatial frequency, isoluminant red/green checkerboard pattern, reversing contrast at 0.5 Hz. The M stimulus was a low contrast (30%), low spatial frequency achromatic checkerboard, counter-phase flickering at 10 Hz. The right side of Figure 1A shows the M and P layers of the LGN from Nissl stained histology (Amunts et al., 2013) and the simulated M-P fMRI pattern (see methods for details about the simulation). Figure 1B shows the group averaged M-P beta maps of the LGN for normal, amblyopia and fellow eye groups. Different eye groups consistently showed a ventral cluster with response bias to the M stimulus, and a dorsal cluster preferred the P stimulus. For each cluster, 30 mm^3^ voxels with the strongest response bias were shown (the highest and lowest M-P beta values for the M and P dominant clusters, respectively). For all three eye groups, the M dominant clusters were located at the ventral LGN, while the P dominant clusters were located at the dorsal LGN. Such spatial pattern was consistent with the simulated M-P pattern from the histology of the human LGN, and was also consistent with the findings from previous studies (Zhang, Zhou, et al., 2015). For the amblyopia eye, response bias to the P stimulus in the P cluster was much weaker compared to the normal and fellow eye groups, suggesting a selective response loss to sustained chromatic stimulus in the P layers of the LGN.

In the ROI analysis, we extracted fMRI responses to the M and P stimuli from the M and P dominant clusters (lower panel of Figure 2B). A leave-one-subject-out procedure was used for the ROI definition to avoid the problem of “double dipping” (Kriegeskorte et al., 2009). For each subject, the M and P ROIs were defined from the average M-P differential beta map of the LGNs of the rest of subjects in the same group. The M dominant cluster was defined as 30 voxels with strongest response bias to the M stimulus, while the P cluster was defined as 30 voxels with strongest preferential response to the P stimulus. Because of individual differences, the leave-one-subject-out ROI definition procedure would reduce the response bias to the M and P stimuli. Thus, the relatively small response bias in the bar graph of Figure 2B was expected. However, a significant response bias to the M and P stimuli were still observed for the normal eye and fellow eye, indicated by a significant interaction between stimuli (M/P stimuli) and layers (M/P layers): F(1,32) = 31.65,*p* < 0.001. Then we compared the LGN responses between the amblyopic eye and normal eye groups. Three-way ANOVA with within-subject factors of stimuli (M/P stimuli) and layers (M/P layers), and between-subject factor of eyes (AE/NE) showed a significant main effect of eyes (F(1,32) = 9.81, *p* = 0.0037), indicating reduced LGN response to the amblyopic eye compared to normal controls. The three-way interaction of eyes × stimuli × layers was also significant (F (1,32) = 10.82, *p* = 0.0024). Two-way ANOVAs showed significant main effect of eyes (F(1,32) = 22.27, *p* < 0.001), and interaction of eyes × stimuli in the P cluster (F(1,32) = 11.32, *p* = 0.002), but not in the M cluster (both*p* > 0.05). Post-hoc t-test showed significant AE response loss to the P and M stimulus in the P cluster *(p* < 0.001 and *p* = 0.0025, respectively), but not in the M cluster (both *p* > 0.05). These findings demonstrate that compared to normal eye’s response, amblyopia eye’s responses to both P and M stimuli were reduced in the P layers but not in the M layers, and there was a stronger response loss to the P stimulus. Comparison between the amblyopic and fellow eyes’ responses showed similar results (main effect of eyes: F(1,16) = 13.15, *p* = 0.0023; interaction of eyes × stimuli × layers: F(1,16) = 4.83, *p* = 0.043; interaction of eyes × stimuli in the P cluster: F(1,16) = 8.52, *p* = 0.01; interaction of eyes × stimuli in the M cluster: F(1,16) = 0.004, *p* = 0.95). Compared to the normal eye, the fellow eye’s response in the LGN showed no significant difference (main effect of eyes: F (1,32) =0.038, p=0.85,*p* > 0.05 for all interactions with the eyes). These findings clearly demonstrated selective reduction of the amblyopia eye’s response to sustained chromatic (P) stimulus in the P layers of the LGN.

At lower magnetic field (<=3T), T2* weighted BOLD response is strongly contributed by non-specific signals from large blood vessels (Gati et al., 1997). According to the vasculature of human LGNs (Abbie, 1933; Schlesinger, 2012), there is a hilar region in the ventral LGN with rich vasculature. Thus, the M layers’ response might be contaminated by non-specific signals from the vascular hilum. However, the dorsal P layers’ response should be less affected. And the differential maps clearly show that there still remains clear response bias (or specific signals) in the M and P dominant clusters. Another possible confounding source to the LGN’s response is the ventral pulvinar, which is closely located to the LGN and is also activated by visual stimuli. However, during data analysis, the LGN voxels were restricted within the anatomical mask of the LGN. In order to further rule out these confounding factors and to replicate our 3T fMRI findings, a 7T fMRI experiment was performed at a higher spatial resolution. Due to short T2* of blood at ultra-high magnetic field, non-specific signals from large draining veins in the vascular hilum should be much reduced, and layer-specific signals from the microvessels (capillaries) will be stronger (Gati et al., 1997; Kim, 2018). With smaller point spread function (PSF) of BOLD signal at 7T and higher spatial resolution, the influence from the ventral pulvinar should be minimal.

#### 1.2 High-resolution 7T fMRI revealed reduced amblyopic eye’s response to high contrast achromatic stimulus in the P layers of the LGN

According to electrophysiological studies, the role of P cells in primate vision are “double duty”: they are not only sensitive to chromatic information, but also responsive to high contrast, high spatial frequency achromatic stimulus (Derrington et al., 1984; Derrington & Lennie, 1984; Hubel & Livingstone, 1990). Thus, if there is a general response deficits to the P cells of the LGN, the amblyopia eye’s response to high spatial frequency luminance-defined stimulus should be significantly reduced in the P layers. To test this hypothesis, and to reduce the influence from non-specific signals as in the 3T fMRI experiment, we took advantage of the resolving power of 7T fMRI to dissociate the M and P layers of the LGN according to their anatomical landmarks. For each voxel in the anatomical ROI of the LGN, we calculated a “layer index” as the normalized Euclidean distance to the dorsal and ventral surfaces of the LGN (ranged from 0 to 1). According the histology of human LGNs, the volume ratio between the M and P layers is about 1 to 4 (Andrews et al., 1997). Based on this M/P volume ratio and the calculated layer indices, we divided the anatomical volume of the LGN into an M and a P layer compartments (right panel of Figure 2A, and also Figure S2).

The visual stimulus was a full contrast, high spatial frequency checkerboard pattern, reversing contrast at 4Hz (Figure 2A, left). According to our recent study (Qian et al., 2020), this achromatic P stimulus should strongly activate the P layers of the human LGN. Indeed, when the fellow eye was stimulated, the achromatic P stimulus strongly activated the dorsal part of the LGN (Figure 2B), corresponding to the P layers. Compared to the fellow eye, P layers’ response to the amblyopic eye was much reduced (as shown by the AE-FE beta map, Figure 2C left). A significant cluster of voxels can be found in the P layer ROI (AE-NE T map, the right most column) at the posterior part of the LGN, corresponding to the central visual field (Connolly & Van Essen, 1984; Schneider et al., 2004). In the ROI analysis (right panel of Figure 2C), permutation test (see methods for details) showed significantly reduced achromatic response in the P layers *(p* = 0.026). Since the achromatic stimulus was designed to selectively activate the P layers of the LGN, the M layers’ response was weak and showed no significant difference between the AE and FE groups *(p* = 0.54). These findings further support amblyopic response deficits to high contrast, high spatial frequency achromatic stimulus in the P layers of the LGN.

#### 1.3 No significant difference of LGN volume between amblyopic patients and normal controls

Anatomical volumes of the LGNs were manually defined from the T1 anatomical images by two experienced researchers blind to the group labels of subjects. All subjects from the 3T and 7T experiments were enrolled in the analysis. The volumes of the left and right LGNs were averaged for each subject. The LGN volume was 132.82 ± 43.20 mm^3^ (Mean ± SD) for normal controls, and 135.26± 42.61 mm^3^ for amblyopia patients. Independent t-test showed no significant difference of LGN volumes between amblyopia patients and normal controls (t(41) = 0.205, *p* = 0.838). Also, the volume of the ipsilateral (temporal retina) and contralateral (nasal retina) LGNs to the amblyopic eye showed no significant difference (F(1, 22) = 0.111, *p* = 0.742, three bilateral amblyopia patients from the 7T data were excluded from this analysis).

### 2. Layer-dependent response loss and enhancement to chromatic P stimulus in the SC

As shown by Figure 3A (upper panel), the M- and P-biased visual stimuli mainly activated the superficial SC, with weaker BOLD response in the deeper part of the SC. This finding is consistent with the consensus that superficial SC mainly processes visual sensory information. The lower panel of Figure 3A show that compared to normal controls, the amblyopia eye’s response to the P stimulus was significantly reduced in the superficial SC, while the fellow eyes response was significantly stronger in the deep SC (likely the intermediate layers). Compared to the fellow eye, the amblyopia eye shows significantly weaker response mainly in the superficial SC.

**Figure 3.**
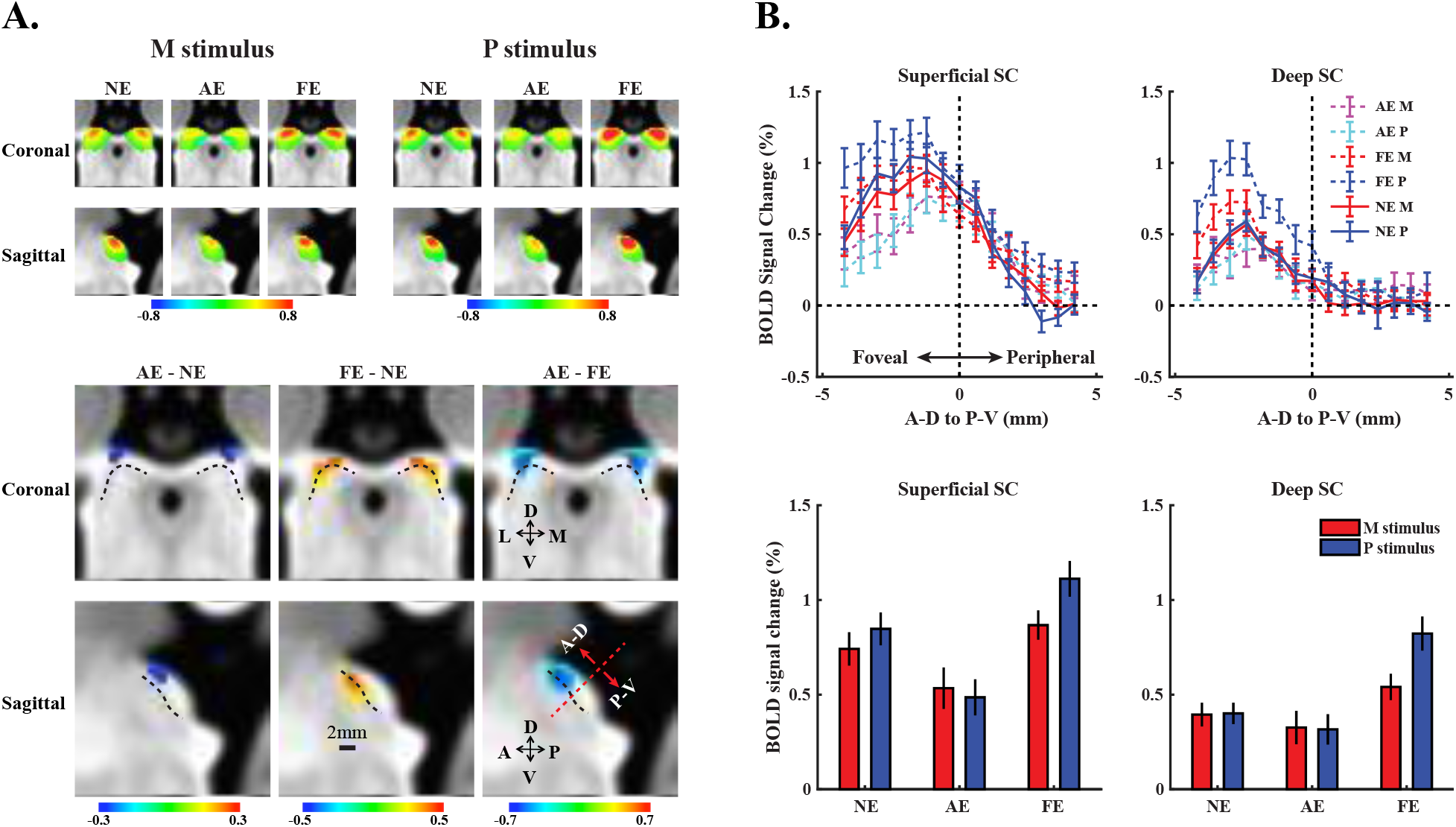
3T fMRI response to the M and P stimuli in the SC. **A. Upper:** Group averaged M and P beta maps for the normal eye (NE), amblyopia eye (AE) and fellow eye (FE) groups, respectively. **Lower:** Voxels with significantly different P responses for AE-NE, FE-AE, and AE-FE comparisons (uncorrected *p* < 0.05, 0.01, 0.001, respectively). Dotted lines indicate the boundary between the superficial and deeper parts of the SC. **B. Upper:** BOLD response in the superficial and Deep SC from anterior-dorsal (A-D, more foveal) to posterior-ventral (P-V, more peripheral) direction. **Lower:** BOLD response in the superficial and deep SC in the anterior-dorsal half of the nucleus, corresponding to the central visual field. Error bars represent standard errors.

During ROI analysis, we first generated a normalized depth map of the SC (from 0 to 1, Figure S3). Since the nominal spatial resolution of the fMRI voxel is at 2mm isotropic which may not be high enough to resolve each individual layers of the SC, and that the functions of the superficial and deeper (middle and deep) layers of the SC is more distinguished in term of visual sensory processing and premotor control and attention, the SC was divided into a superficial and a deeper part (at normalized depth = 0.5). In Figure 3B (upper panel), we plotted BOLD responses in the superficial and deep SC from anterior-dorsal to posterior-ventral direction. Results show that strongest responses were located at the anterior-dorsal part of the nucleus, corresponding to the central visual field. Thus, we calculated statistics only from the anterior-dorsal part of the SC. First, we compared the responses between the amblyopic and normal eye groups. Three-way ANOVA of eyes (AE/NE), stimuli (M/P), and layers (superficial/deep) showed a significant interaction of eyes × layers (F(1,32) = 4.14, *p* = 0.05), and a significant interaction of eyes × stimuli × layers (F(1,32) = 4.87, *p* = 0.035). Two-way ANOVA showed that the eyes × layers interaction was significant for the P stimulus (F(1,32) = 6.7, *p* = 0.014), but not for the M stimulus (F(1,32) = 1.78, *p* = 0.19). Post-hoc t-test showed significant difference between the AE and NE response to the P stimulus in the superficial SC (t(32) = 2.2, *p* = 0.036), but not in the deep SC (t(32) = 0.69, *p* = 0.49). These results indicate that compared to normal controls, amblyopia eye’s response to the chromatic P stimulus was selectively reduced in the superficial but not in the deep SC. Then we compared the responses between the fellow and normal eye groups. Three-way

ANOVA of the FE and NE response showed a significant interaction of eyes × stimuli (F(1,32) = 5.28, *p* = 0.028), and significant interaction of eyes × stimuli × layers (F(1,32) = 4.63, *p* = 0.039). Two-way ANOVA showed significant main effect of eye (F(1,32) = 4.92, *p* = 0.034), and interaction of eyes × stimuli in the deep SC (F(1,32) = 14.04, *p* < 0.001), but not in the superficial SC (both*p* > 0.2). Post-hoc t-test showed significantly stronger FE response in the deep SC to the P stimulus (t(32) = 3.1, *p* = 0.004), but not to the M stimulus *(p* = 0.16). These results indicate that compared to normal controls, the fellow eye’s response to the chromatic P stimulus was selectively stronger in the deep SC. Finally, for the comparison between the amblyopic and fellow eyes, three-way ANOVA showed significant main effect of eyes (F(1,32) = 6.29, *p* = 0.023), and interaction of eyes × stimuli (F(1,32) = 13.97, *p* = 0.002), but the three-way interaction was not significant (F(1,32) = 0.035, *p* = 0.85). It indicates that compared to the fellow eye, the amblyopia eye’s response to the P stimulus was significantly weaker in both superficial and deep SC.

### 3. Selective amblyopic response loss to chromatic P stimulus in the visual pulvinar

In Figure 4, the M and P visual stimuli significantly activated the inferior portion of the pulvinar, corresponding to the visual part of the nucleus (Arcaro et al., 2015; DeSimone et al., 2015). According to the human subcortical brain atlas from Nissl stained histology (Mai et al., 2015), both the ventral and the dorsal parts of the inferior pulvinar were activated. For the normal and fellow eye groups, BOLD responses in the visual pulvinar were strongly biased to the P stimulus, while the amblyopia eye’s response showed much weaker response to the P stimulus. During ROI analysis, a leave-one-subject-out procedure was used to define the ROI of visual pulvinar significantly activated by the M and P stimuli (see methods for details). The bar plot shows that the amblyopic eye’s response to the P stimulus was strongly reduced, but the response to the M stimulus was almost unaffected. Two-way ANOVA of eyes (AE/NE) and stimuli (M/P) showed significant main effect of eye (F(1,32) = 5.08, *p* = 0.031), and interaction of eyes × stimuli (F(1,32) = 13.27,*p* < 0.001). Post-hoc t-test showed significantly reduced AE response to the P stimulus (t(32) = 3.43, *p* = 0.0017), but not to the M stimulus *(p* = 0.559). Twoway ANOVA of AE and FE responses showed similar results, with significant interaction of eyes × stimuli (F(1,16) = 18.78, *p* < 0.001), and post-hoc t-test shows significant difference for the P response but not for the M response *(p* = 0.021 and 0.579, respectively). No significant difference was found between the FE and NE responses (all *p* > 0.4).

**Figure 4.**
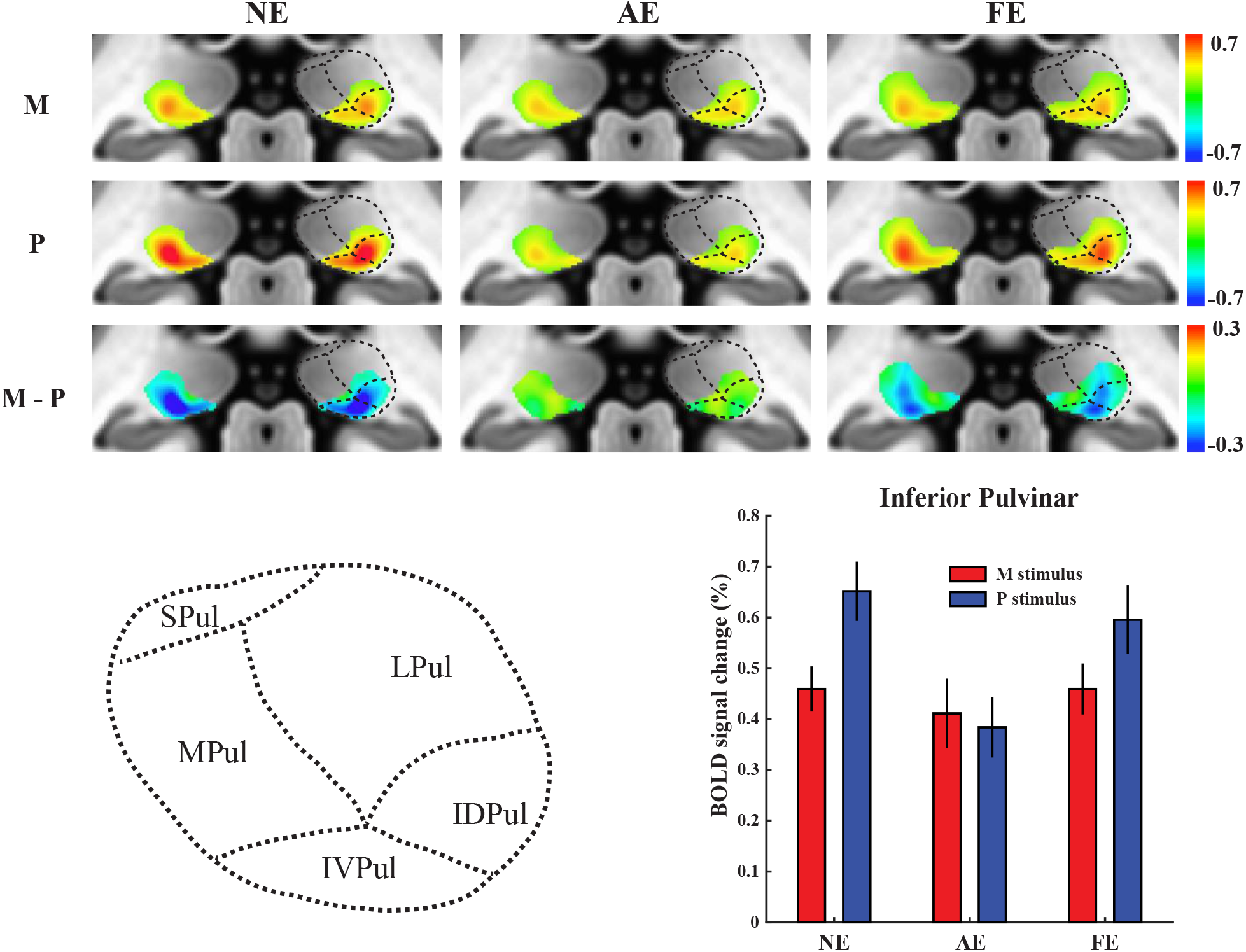
3T fMRI responses to the M and P stimuli in the visual pulvinar. **Upper:** Group averaged M-P beta maps of visual pulvinar (MNI y = 30mm). Maps were thresholded at M+P*p* < 0.001 uncorrected. **Lower left:** Subregions of the pulvinar according the human brain atlas based on Nissl stained histology (Mai, Majtanik et al. 2015). Black dotted lines indicate the approximate boundaries between the subregions. **Lower right:** Responses in the ROI of the inferior pulvinar. Error bars represent standard errors.

### 4. Selective amblyopic response loss to chromatic stimulus in the early visual cortex (V1-V3) and ventral visual stream (V4/VO/LO)

Figure 5 shows the normalized fMRI responses to the amblyopic and fellow eyes (divided by the response to the control eyes) in the subcortical and cortical visual areas. A subject-level bootstrapping procedure was used for the normalization of fMRI response (see method for details). Normalized response to the amblyopic eye was more reduced to the P stimulus than to the M stimulus in the early visual cortex (V1-V3), and in the ventral visual areas (V4/VO/LO), but not in the dorsal visual areas (V3ab/MT). Notably, amblyopic loss of parvocellular response was much less in the cortical areas compared to that in the LGN (LGNp vs. V1: p < 0.001), suggesting that amblyopic deficit in the LGN may not be entirely inherited from cortical feedbacks. Normalized response to the fellow eye showed no significant difference from normal controls in most cortical areas. However, the fellow eye’s response to the P stimulus in the deep SC was significantly stronger. Figure S4 shows the original BOLD responses in the visual cortex to the M and P stimuli. We also examined the difference of responses from anisometropia and strabismus patient groups. A number of visual areas showed significant difference, especially in the higher order visual cortex (Figure S5).

**Figure 5.**
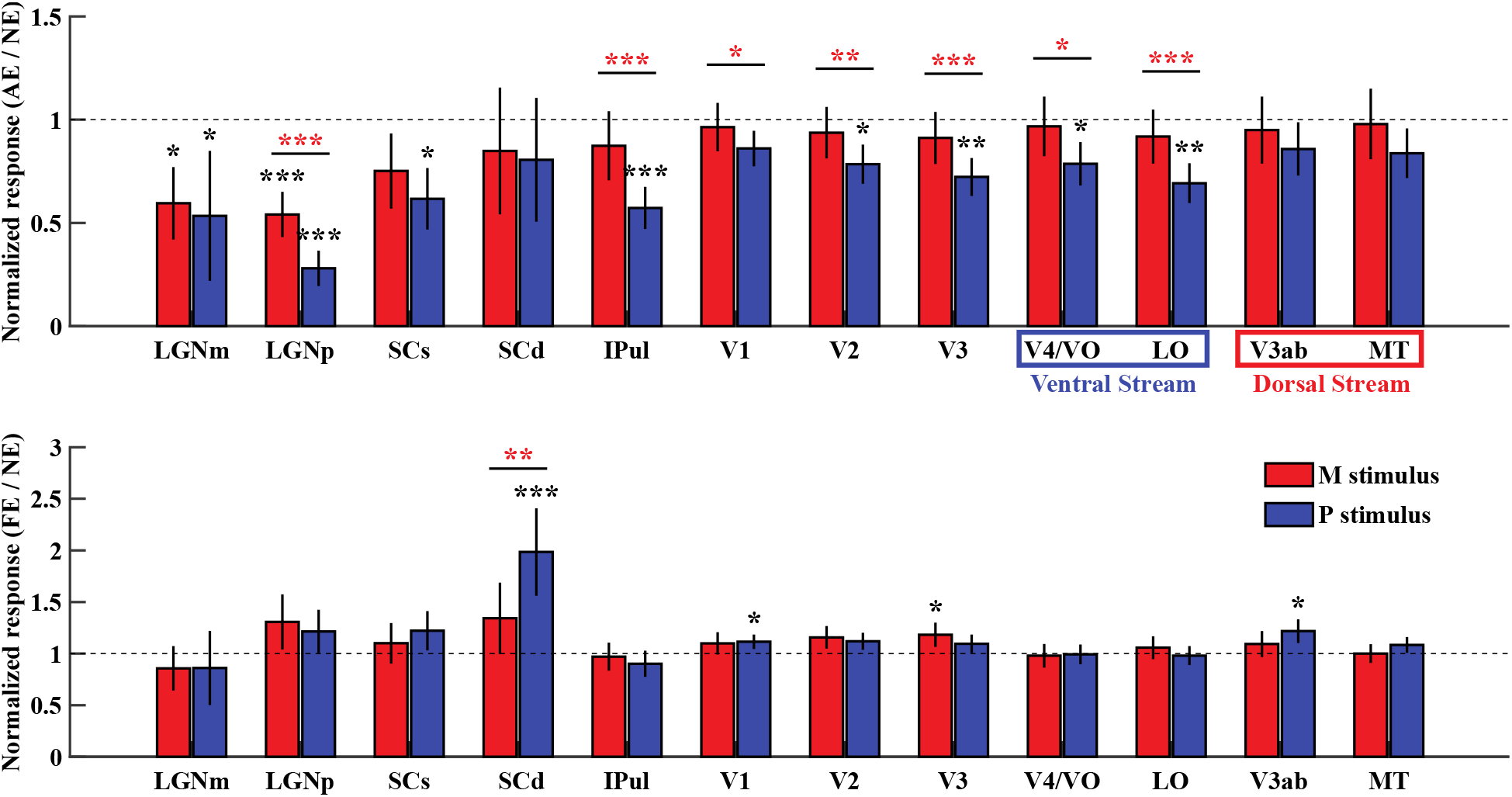
Normalized fMRI response to the amblyopic and fellow eyes in the subcortical nuclei and cortical visual areas. Upper and lower panels show the normalized response to the amblyopic eye and Γello* eye, respectively (divided by the response of normal controls). Essor bars Sodicate standard deviation of the bootstrapped distribution of the mean. Abbreviations, LGNm: M layers of the LGN, LGNp: P layers of the LGN, IPul: Inferior Pulvinar, SCs: superficial SC, SCd: deep SC. *\*p* < 0.05, *\*\*p* < 0.01, *\*\*\*p* < 0.001.

### 5. Dynamic causal modeling of effective connectivity

To understand the cause of amblyopic deficit to subcortical nuclei, we used Dynamic Causal Modeling (DCM) to investigate the effective connectivity of fMRI signals between visual areas, and the connectivity difference between different eye groups (Friston et al., 2003). As shown by Figure 6 (upper row), we first defined a full model based on known anatomical connections of the primate visual system (Briggs & Usrey, 2011; Felleman & Van Essen, 1991; May, 2006; Sherman & Guillery, 2002; Shipp, 2003; Ungerleider et al., 1984). In this full model, the LGN receives both M and P driving inputs from the retina, while the SC only receives the M driving input. Fixed connections were defined between and within visual areas, which could be modulated when the M or P stimuli were presented. Connectivity strength of the full DCM model was estimated for each individual (Zeidman, Jafarian, Corbin, et al., 2019). At the group level, we used Bayesian model reduction, Bayesian model average and Parametric Empirical Bayes to make inference about the connectivity strength and the differences between eye groups (Friston et al., 2016; Zeidman, Jafarian, Seghier, et al., 2019).

**Figure 6.**
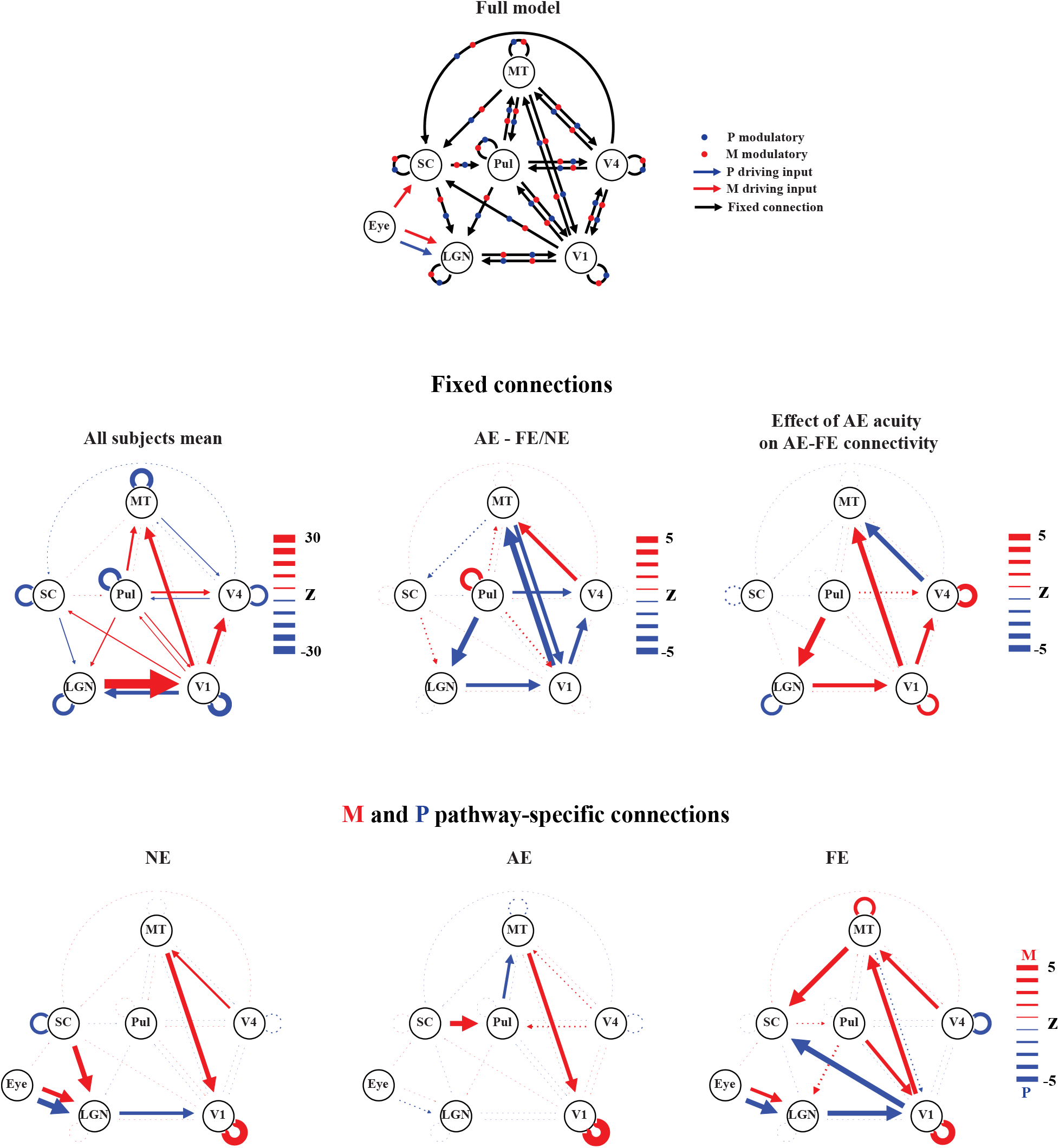
Dynamic causal modeling of effective connectivity. The upper row shows the full DCM model. Figures from the middle row show the results for the fixed connections. Red and blue arrows denote positive and negative connections, respectively. Lower panels show the M and P pathway-specific connections, including the M and P driving input to the LGN and SC, and the difference between the M (red) and P (blue) modulatory connections. Line width and arrow size represent Z value of the connection. Solid and dashed lines represent significant *(p* < 0.05) and non-significant connections *(p* > 0.05), respectively.

The mean of fixed connections across all subjects showed robust feedforward connectivity from the LGN to V1 (Figure 6, left panel of the middle row), which was significantly reduced for the amblyopic eye compared to the normal and fellow eyes (Figure 6, middle panel of the middle row). Fixed connections from pulvinar to the LGN and pulvinar to V4, as well as several cortico-cortical connections were also significantly reduced for the amblyopic eye. Compared to the fellow eye, stronger LGN → V1 and pulvinar → LGN connectivity driven by the amblyopic eye predict better visual acuity of the amblyopic eye (Figure 6, right panel of the middle row). The lower row of figure 6 shows the M and P pathway-specific connections, which is the difference between the M and P modulatory connectivity. Compared to the M stimulus, P stimulus presented to the normal and fellow eyes significantly increased the LGN → V1 connectivity. This P-pathway specific feedforward connectivity was not significant driven by the amblyopic eye’s input. The M and P driving inputs to the LGN were also significantly reduced for the amblyopic eye compared to the normal and fellow eyes (Bootstrap test: *p* = 0.03 and *p* = 0.027 for the M and P driving inputs, respectively), suggesting reduced feedforward processing of the LGN. For the fellow eye, there were significant parvocellular feedback from V1 and magnocellular feedback from MT (right panel in the lower row) to the SC, which was absent in the normal eye and amblyopic eye groups. It suggests that increased fellow eyes’ response in the deep SC might be due to cortical feedbacks. Amblyopic eye also showed significant magnocellular connectivity from the SC to pulvinar, which may suggest intact feedforward processing of the tectal-pulvinar pathway.

### 6. Reduced LGN activity cannot be explained by fixation instability of the amblyopic eye

Amblyopic eyes, especially those with strabismus, showed unsteady fixational eye movements, such as higher frequency, amplitude and speed of microsaccades, and higher amplitude and speed of slow drifts (Chung et al., 2015). To evaluate the potential effect of fixation instability on the BOLD response of early visual areas, we performed an additional fMRI experiment on normal subjects viewing stimuli monocularly with and without artificially imposed jerky movements to simulate the unsteady fixation of strabismus eyes. In the normal eye viewing condition, subjects viewed the same M and P stimuli as in figure 1A; in the strabismus eye viewing condition, jerky motion was introduced to the visual stimuli to simulate the unsteady fixational eye movements of strabismus eyes. Figure S5A shows the example trajectories of simulated eye movements of the strabismus eye. Characteristics of simulated eye movements largely matched the statistics of strabismus eye movements from a recent study (Chung et al., 2015): area of 68% bivariate contour ellipse (BCEA, 0.21 deg^2^), microsaccade frequency (2.9 Hz), microsaccade amplitude (26.9 arc min), slow drift speed (2.64 deg/sec), slow drift amplitude (11.2 arc min). During the experiment, subjects were instructed to maintain fixation while passively viewing the visual stimuli. Two-way repeated-measures ANOVA with within-subject factors of stimuli (M/P stimuli) and eyes (normal/strabismus eyes) showed no significant main effect of eyes on the BOLD response of the LGN (F(1,7) = 0.35, *p* = 0.573) and V1(F(1,7) = 0.896, *p* = 0.375). Post-hoc t-test also showed no significant difference between the normal and strabismus eye viewing conditions for either M or P stimuli *(p* > 0.05 for all comparisons). These results suggest that reduced BOLD activity of the LGN was unlikely due to fixation instability of the amblyopic eye.

## Discussion

Using stimuli to preferentially activate the M and P layers of the LGN, we found that amblyopia led to selective response loss to sustained chromatic stimulus in the P layers of the LGN. High-resolution 7T fMRI further demonstrated response loss to high contrast, high spatial frequency luminance-defined stimulus in the P layers, but not in the M layers of the LGN. Feedforward connectivity from the LGN to V1 was also significantly decreased, which correlated with the visual acuity loss of the amblyopic eye. These findings demonstrated amblyopic functional deficits on feedforward processing in the P layers of the LGN. Amblyopic deficits were also observed for parvocellular responses in the visual pulvinar, early visual cortex, and the ventral but not the dorsal visual streams. The effective connectivity from the pulvinar to V4 also significantly decreased. Another key finding is that the amblyopic eye’s response to the chromatic stimulus was reduced in the superficial SC; whereas the fellow eye’s response robustly increased in the deeper layers of the SC, associated with pathway-specific feedbacks from the visual cortex. It is worth to mention that our patients had not received prior treatment for amblyopia, thus these subcortical functional abnormalities should be results from nature history.

Previous psychophysics and neuroimaging studies investigated the M and P pathway-specific functional deficits of amblyopia. However, lacking a non-invasive tool to directly access layer-specific responses of the human LGN, these indirect evidences are controversial. Levi and Harwerth found contrast sensitivity loss for amblyopia eye most pronounced at high spatial frequencies (Levi & Harwerth, 1977). However, using a luminance pedestal discrimination paradigm, Zele and colleagues found that contrast sensitivity was similarly affected for M and P biased stimuli (A J Zele et al., 2007; Andrew J. Zele et al., 2010). Some neuroimaging studies of human visual cortex found reduced cortical response to high spatial and low temporal frequency stimuli (Barnes et al., 2001; Choi et al., 2001; Lerner et al., 2006), while others showed selective response loss to low spatial and high temporal stimuli (Hess, Li, et al., 2009; Li et al., 2013). A recent fMRI study showed more reduced response to chromatic stimulus than to achromatic stimulus in the LGN of amblyopia patients (Hess, Thompson, et al., 2010), but the M and P layer-specific responses were not distinguished. Since the P cells respond well to both chromatic and achromatic stimuli, it is unclear whether this result was due to selective response loss to chromatic than to achromatic stimuli in the P layers, or due to selective loss to chromatic response in the P layers than to achromatic response in the M layers of the LGN. Moreover, the spatial resolution of fMRI in this study was quite low, thus the BOLD response of the LGN was very likely to be contaminated by the response of visual pulvinar which locates closely to the LGN. In the current study, the high-resolution fMRI approaches minimized the contamination of BOLD signal from the visual pulvinar. Our 3T and 7T fMRI results showed clear amblyopic loss to both chromatic and achromatic responses in the P layers of the LGN, but less reduction to achromatic response in the M layers. Effective connectivity analysis revealed reduced feedforward connectivity from the LGN to V1, which is significantly correlated with the acuity loss of the amblyopic eye. It is consistent with previous studies showing structural abnormalities of optic radiation in adult and children with amblyopia (Allen et al., 2015; Qi et al., 2016; Xie et al., 2007). These findings provide direct and strong evidence for selective loss of feedforward processing in the P layers of the LGN of adult human amblyopia.

Visual pulvinar receives driving input from the visual cortex, and regulates information transmission across cortical areas (Purushothaman et al., 2012; Zhou et al., 2016). It also receives input from the SC and sends projections to extra-striate and posterior parietal cortex (Adams et al., 2000; Berman & Wurtz, 2010; Lyon et al., 2010). The influence of amblyopia to the function of pulvinar remains almost unknown. Only one previous study showed minimal amblyopic loss of motion response in the pulvinar (Thompson et al., 2012). In the current study, we found very robust amblyopic response loss to the sustained chromatic (P) stimulus in the inferior pulvinar, while the response to the transient achromatic (M) stimulus was almost unaffected. Effective connectivity from pulvinar to V4 was also significantly reduced for stimuli presented to the amblyopic eye. These findings demonstrate parvocellular response deficits in the visual pulvinar and abnormal information transmission through the pulvino-cortical pathway. However, amblyopic eyes’ input produced significant magnocellular feedforward connectivity from the SC to ventral pulvinar, suggesting intact magnocellular processing through tectal-pulvinar pathway.

To the best of our knowledge, the influence of amblyopia to the function of retinotectal pathway has never been investigated in primate. A previous study on rodent found reduced glucose metabolism in the superficial layers of SC by monocular lid suture during the critical period (Wang et al., 2005). Our human fMRI results clearly demonstrated that the amblyopic eye’s response in the superficial SC was significantly more reduced to sustained chromatic P stimulus than to transient achromatic M stimulus. Since the superficial SC does not receive direct projection from the midget (P) ganglion cells of the retina, such chromatic response deficit should be related to feedbacks from the visual cortex (Finlay et al., 1976; White et al., 2009). According to computational models and recent empirical studies (White, Berg, et al., 2017; White, Kan, et al., 2017; Yan et al., 2018; Zhaoping, 2008), cortical feedbacks to the superficial SC might be related to the representation of an attention saliency map. Thus, reduced chromatic response in the superficial SC may indicate a degraded representation of the attention saliency map. An interesting and surprising finding is that compared to control eyes, the fellow eye’s response to the chromatic P stimulus strongly increased in the deep SC, associated with a significant parvocellular feedback connectivity from V1 to the SC. V1 response to the P stimulus presented the fellow eye also significantly increased (Figure 5). These findings suggest that increased fellow eyes’ response in the deep SC should be due to cortical feedbacks. Although we didn’t record eye movements in the scanner, several previous studies found dysfunctions of fixational drifts and microsaccades of the amblyopia eye but not of the fellow eye during monocular viewing (Chung et al., 2015; Gonzalez et al., 2012; Shaikh et al., 2016; Subramanian et al., 2013). However, it might be possible that the fellow eye enhanced fixational eye movements (e.g. more microsaccades) specifically to stimulus with more spatial details, such as the high spatial frequency P stimulus used in the current study. Improved miniature eye movements of the fellow eye might help better sampling of fine spatial details to compensate acuity loss of the amblyopia eye (Kuang et al., 2012; Rucci et al., 2007). Another related explanation is a compensation mechanism of attention in the SC boosting the fellow eye’s response. Future studies should link the psychophysics and neuroimaging data to understand the functional significance of SC response changes in amblyopic vision.

To summarize, the current study demonstrates that abnormal visual experience during critical period shifts the homeostatic interocular balance of human subcortex in a pathway-specific manner. Amblyopia led to selective deficit of parvocellular feedforward signals in the geniculostriate pathway, whereas loss and compensation of parvocellular feedback signals in the retinotectal pathway. These pathway-specific functional abnormalities provide the neural basis for the loss of visual acuity, and deficits in control of eye movement and attention in amblyopia. It sheds lights for developing new tools for effective and precision treatment of the disease, such as perceptual learning to selectively improve the amblyopic eye’s sensitivity in the parvocellular geniculostriate pathway, or eye-specific training of attention targeting the retinotectal pathway. Our high-resolution fMRI approach may serve as direct and quantifiable biomarkers for diagnosis and tracking the prognosis of amblyopia, in addition to traditional behavioral measures of ophthalmology.

## Materials and Methods

### Subjects

17 adult patients diagnosed with unilateral amblyopia (10 anisometropia and 7 strabismus patients, age 25.0±5.0 years, 4 females) and 17 gender matched healthy controls (age 33.0±5.6 years) were enrolled in 3T fMRI study. Previous studies showed that BOLD response is greatly reduced in early visual areas of amblyopia (Hess, Li, et al., 2010; Hess, Thompson, et al., 2009) (Cohen d ≫ 1 from both studies). The sample size (N=17 for both groups) provides at least 80% power to detect a large difference between two groups (Cohen d > 1). 9 adult patients diagnosed with anisometropia (6 unilateral and 3 bilateral patients, age 29.78±8.21 years, 6 females) were enrolled in the 7T fMRI study. The sample size (N=9) provides at least 75% power to detect a large within-subject difference between amblyopic and fellow eyes (Cohen d > 1). The research followed the tenets of the Declaration of Helsinki, and all participants gave written informed consent in accordance with procedures and protocols approved by the Human Subjects Review Committee of the Eye and ENT Hospital of Fudan University, Shanghai, China. 7T participants also gave written informed consent approved by institutional review board of the Institute of Biophysics, Chinese Academy of Sciences.

The best-corrected visual acuity (BCVA) of each subject was assessed by experienced optometrist with the Snellen chart. Cover and alternative cover testing, duction and version testing, intraocular pressure testing, slit-lamp testing, indirect fundus examination after pupil dilation, and optometry were performed for each subject. The BCVAs of all amblyopic eyes was 20/30 or worse, and the BCVA of all fellow eyes was 20/20 or better. Both amblyopia and control participants were required to be free from a history of intraocular surgery, any eye diseases, and any systemic diseases known to affect visual function (e.g., migraine, congenital color deficiencies). None of amblyopia subjects underwent amblyopia treatment including patching treatment. All strabismic amblyopia patients had undergone strabismus surgery at least one year previously and had normal eye position and steady fixation. Eye dominance in normal subjects was determined by instructing the subject to look at a distant letter through a hole between her/his hands with their eyes alternatively closed and to report the target was visible or invisible. Dominant eye was the eye that still could maintain the letter in the hole (hole-in-card method). Table 1 shows the clinical characteristics of amblyopia patients. Table S1 shows the details for each individual.

**Table 1.** Clinical characteristics of amblyopia patients

| Characteristics | Total patients | Anisometropic | Strabismic |
| --- | --- | --- | --- |
| Age (y, Mean±SD) | 23.76±6.08 | 24.10±6.64 | 23.29±3.25 |
| Gender (percent female) | 41.17% | 40.00% | 42.85% |
| BCVA (Snellen Chart) | 0.362±0.214 | 0.465±0.180 | 0.214±0.175 |
| Eye alignment | -- | Ortho | all esotropia (+14.29° ±5.35°) |

### Stimuli and procedures for the 3T fMRI experiment

Visual stimuli were generated in MATLAB (Mathworks Inc.) with Psychophysics Toolbox extensions (Brainard, 1997; Pelli, 1997), running on a MacBook pro computer, and presented on a translucent screen with an MRI compatible projector at a resolution of 1024 x 768 @ 60Hz. Color lookup table of the display was calibrated to have linear luminance output.

As shown by Figure 1A, full-field (24 degrees in diameter) M and P biased visual stimuli differed in spatiotemporal frequency, contrast and chromaticity were used to selectively activate the M and P layers of the LGN. The M stimulus was a low-spatial-frequency sine wave checkerboard counter-phase flickering at 10Hz with 30% luminance contrast. The P stimulus was a high-spatial-frequency, isoluminant red/green square wave checkerboard reversing contrast at 0.5Hz. The P stimulus rotated clockwise or counterclockwise at 2.4 degrees/seconds to prevent adaptation. The subjective isoluminance of red and green was adjusted with a minimal flicker procedure, repeated four times for each subject in the scanner before the fMRI experiment. To identify the region of interest (ROI) of the LGN of visual pulvinar, hemifield checkerboard stimuli counter-phase flickering at 7.5 Hz were presented in-alternation to the left and right visual field every 16 seconds.

The fellow eye and the amblyopic eye of amblyopia patients were tested in separate sessions. Each session consisted of four runs for the M and P visual stimuli, two runs for the hemifield LGN localizer, and a T1 anatomical scan. Each fMRI run lasted 256 seconds, with two dummy scans before each run. The non-dominant eye of the normal controls was tested with the same experimental procedure. During fMRI scanning, subjects were instructed to maintain fixation and pay attention to the visual stimuli. A black eye patch covered the non-test eye, and the test eye wore glasses to correct refractive errors.

### Stimuli and procedures for the 7T fMRI experiment

Full contrast checkerboard stimuli counter-phase flickering at 4Hz was used and the diameter is 10 degree (Figure 2A). The left and right eyes of amblyopia patients were tested in the same scanning session and each session consisted of six runs: three runs for the left eye and anther three for the right eye. Each functional run lasted 276 seconds, with 16 seconds of visual stimulation (8 blocks) interleaved with 16 seconds of fixation period (9 blocks). Two dummy scans were collected in the beginning of each run and were discarded during data analysis. During the experiment, the subjects were instructed to maintain fixation and to pay attention to the stimulus. Both eyes wore glasses to correct refractive errors. A black eye patch covered the non-tested eye. After one eye finished scanning, subject switched the eye patch in the scanner by themselves.

### MRI Data acquisition

3T MRI data were collected with a 3T scanner (Siemens, Verio) at the Eye and ENT Hospital of Fudan University using a twelve-element head matrix coil. A reduced field of view (FOV) gradient echo planar imaging sequence was used to acquire functional images (2mm isotropic voxels, 26 axial slices, 64×64 matrix with 2mm in-plane resolution, 2mm thickness, TR/TE = 2000/28 ms, flip angle = 90°, a saturation band was placed on the frontal regions to avoid image wrap-around). Reducing the FOV significantly reduces the echo train length of EPI, thus alleviates image distortion and blur of high resolution imaging (Heidemann et al., 2012). Anatomical volume was obtained with a T1 MPRAGE sequence (1mm isotropic voxels, 192 sagittal 1mm-thick slices, 256 × 256 matrix with 1mm in-plane resolution, TR/TE = 2600/3.02 ms, flip angle = 8°).

7T fMRI data were acquired on a 7T MRI scanner (Siemens Magnetom) with a 32- channel receive 1-channel transmit head coil (Nova Medical), from Beijing MRI center for Brain Research (BMCBR). Gradient coil has a maximum amplitude of 70mT/m, 200us minimum gradient rise time, and 200T/m/s maximum slew rate. Functional data were collected with a T2*-weighted 2D GE-EPI sequence (1.2mm isotropic voxels, 26 axial slices with 1.2mm slice thickness, TR = 2000 ms, TE = 22 ms, image matrix =150×150, FOV = 180×180 mm, GRAPPA acceleration factor = 2, Flip angle=80°, partial Fourier = 6/8, phase encoding direction from A to P). Before each functional scan, five EPI images with reversed phase encoding direction (P to A) were also acquired for EPI distortion correction. Anatomical volumes were acquired with a T1-MP2RAGE sequences at 0.7 mm isotropic resolution (256 sagittal slices, centric phase encoding, acquisition matrix=320×320, Field of view = 224×224 mm, GRAPPA=3, TR = 4000 ms, TE=3.05 ms, TI1 = 750ms, flip angle = 4°, TI2 = 2500 ms, flip angle = 5°). Subjects used bite bars to restrict head motion.

### MRI Data analysis

#### Preprocessing

Preprocessing of functional data was done in AFNI (Cox, 1996), including the following steps: physiological noise removal with retrospective image correction, slice timing correction, EPI image distortion correction with nonlinear warping (PE blip-up, for 7T data only), rigid body motion correction, alignment of corrected EPI images to T1-weighted anatomical volume (cost function: lpc), and per run scaling as percent signal change. To minimize image blur, all spatial transformations were combined together and applied to the functional images in one resampling step. General linear models with a fixed HRF (Block4 in AFNI) were used to estimate BOLD signal change from baseline for each stimulus condition. Motion parameters were included as regressors of no interest. Subcortical data were transformed to MNI space using advanced normalization tools (ANTs) (Avants et al., 2011). We used the symmetric MNI template from CIT168 atlas for nonlinear warping (Pauli et al., 2018). Functional data of the subcortex were resampled to 0.6mm isotropic in MNI space. MNI space data were used for the ROI analysis of the SC and Pulvinar, native space data (upsampled to 1mm, and 0.6mm isotropic for the 3T and 7T data, respectively) were used for the LGN ROI analysis.

#### ROI definition and analysis

##### LGN

For the 3T experiment, ROIs of the LGN were defined as voxels significantly activated by the hemifield localizer (uncorrected *p* < 0.05) within the anatomical mask of the LGN. Anatomical ROIs of the LGN were defined by two experienced researchers from T1-MPRAGE images, in which the LGN appeared darker than surrounding tissues. Care was taken not to include activations from the inferior pulvinar. Beta values of the M and P stimuli were extracted from all voxels of the ROI and entered group-level analysis in MATLAB. For each eye group, the coordinates of the left LGNs were mirror-flipped to the right, and all LGNs were registered to the center of mass location. A median LGN mask was then generated (voxel has values from half of the LGNs) as the registration template. Then a twelve-parameters linear (affine) transformation to the median LGN template was performed on each LGN. This procedure ensured approximate matching of orientation and size of the LGNs. Then group level analysis was performed on this “standard space”. M and P ROIs were defined from the M-P beta map. In order to avoid the problem of ‘double dipping’, we used a leave-one-subject-out procedure to define the M and P layer ROIs from the M-P beta map. For each subject, the M and P biased ROIs were generated from the M-P beta map of the rest of subjects in the same eye group. The M ROI was identified as 30 voxels with strongest response bias to the M stimulus (highest M-P values), while the P ROI was defined as 30 voxels with strongest response bias to the P stimulus (lowest M-P values). ROI of the LGNs for the 7T experiment were defined only from the T1-weighted anatomical volumes. The ROI registration procedure was similar to the 3T experiment, except that an LGN template from the MNI space was used. A normalized layer index map was generated on the LGN template. Two layers of voxels were defined corresponding to the ventral and dorsal surfaces of the LGN. For the rest of voxels, we calculated a layer index as the normalized distances to the dorsal surface (ranged from 0 to 1). The ROIs for the M and P layers were determined from the layer index map according the volume ratio of M/P layers of human LGNs (M/P = 1/4) (Andrews et al., 1997).

##### SC

ROI analysis for the SC was performed with MNI space data from the 3T experiment. ROIs of the SC were defined manually from the CIT168 MNI T1 template. To generate the depth map of the SC, two layers of voxels were first defined from the superficial and deep surface of the SC. Then a normalized depth map was calculated for each voxel as the ratio of shortest distances to the superficial and deep surfaces (from 0/superficial to 1/deep). The volume of the SC was split into a superficial and a deeper part at depth = 0.5. To generate the AD to PV slice profile in Figure 3B, the peak response in the superficial and deep ROIs were selected for each slice.

##### Pulvinar

ROI analysis for the pulvinar was also performed with the MNI space data from 3T. Anatomical ROIs of the Pulvinar were defined from the CIT168 MNI template. Spatial smoothing was performed within the ROI with a 1.2mm FWHM Gaussian filter. For each subject, a leave-one-subject-out-procedure was used to define the functional ROI, 300mm^3^ voxels with the strongest M+P beta values from the rest subjects in the group were selected as the ROI.

##### Visual cortex

Retinotopic ROIs of cortical visual areas were defined on the cortical surface according to the polar angle atlas from the 7T retinotopic dataset of Human Connectome Project (Benson et al., 2018).

##### Statistics

A subject-level bootstrapping procedure was used to calculate the statistics of 7T response in Figure 2C (right), the normalized fMRI response in Figure 5, and test the difference in driving input strength between the AE and NE/FE groups (Figure 6). For each permutation, the data for a group of subjects were resampled with replacement. Then the group averaged fMRI responses for different eye groups were calculated. The bootstrapping procedure was repeated 10^6^ times. A null distribution was generated by shifting the mean of bootstrapped distribution to zero, and a two-sided p value was derived from the null distribution.

#### Dynamic causal modeling

Effective connectivity of fMRI data were analyzed using the DCM module of SPM12 (version 2020-Jan-13th). FMRI datasets were preprocessed in AFNI with slice timing correction and motion registration. The mean time course of the LGN was averaged from the ROI of the whole LGN (voxels significantly activated by the hemifield localizer, within the anatomical mask of the LGN). For the SC, pulvinar, V1, MT and V4, time courses from one hundred most significantly activated voxels in the ROI (sorted according to the F value of M+P contrast) were extracted and averaged. For each session, the full DCM model was estimated separately for each run and then averaged using Bayesian fixed effect (FFX) averaging method (spm_dcm_average). The second (group) level analysis used the Parametric Empirical Bayes (PEB) module of SPM (spm_dcm_peb and spm_dcm_peb_bmc). For the pathway non-specific (fixed) connections, a second-level design matrix was defined for commonality (mean across all subjects) and eye effect (AE – FE/NE). Another PEB analysis was performed on the modulatory effect of amblyopia (versus the fellow eye) on connectivity, commonality and the visual acuity of amblyopic eyes were used as the second level regressors. For the pathway-specific connections, a within-subject design matrix was defined as the difference between the M and P modulatory connections, and a between-subject design matrix was defined for the commonality (mean across subjects) separately for different eye groups. Z value of the connection was defined as the mean divided by the deviation of second level parameters. P values of the comparison of driving inputs to the LGN between AE and FE/NE groups were determined by bootstrap hypothesis test.

## Acknowledgements

This study was funded by Beijing Science and Technology Q2 Project (grant nos. Z181100001518002); Strategy Priority Research Program of Chinese Academy of Science, Grand No. XDB32020200; Bureau of International Cooperation, Chinese Academy of Sciences (grant no. 153311KYSB20160030) and National Natural Science Foundation of China (grant no. 31871107, 31930053, 81500752, 81525006, 81670864 and 81730025); Excellent Academic Leaders of Shanghai (18XD1401000).

## Supplemental Materials

### M-P pattern simulation for the 3T BOLD fMRI experiment

As shown by Figure S1, we first manually identified each individual layer of the LGN from Nissl stained brain slices at 20μm resolution (Amunts, Lepage et al. 2013). The neural image for each stimulus was generated by labeling in the corresponding layers with a normalized response magnitude: 1 for the P layers, and 4 for the M layers, based on much higher sensitivity of the M cells (Wiesel and Hubel 1966, Derrington and Lennie 1984). Then spatial smoothing with a 3.5mm FWHM gaussian filter was applied on the neural pattern to simulate the blurring effect of GE BOLD point spread function at 3T (Engel, Glover et al. 1997). Then the BOLD image was resampled to the resolution of fMRI measurement (2mm isotropic) and up-sampled to 0.6mm isotropic with cubic interpolation as in the fMRI data analysis, finally a differential image was generated by subtracting the patterns between the M and P stimulus conditions.

**Figure S1.**
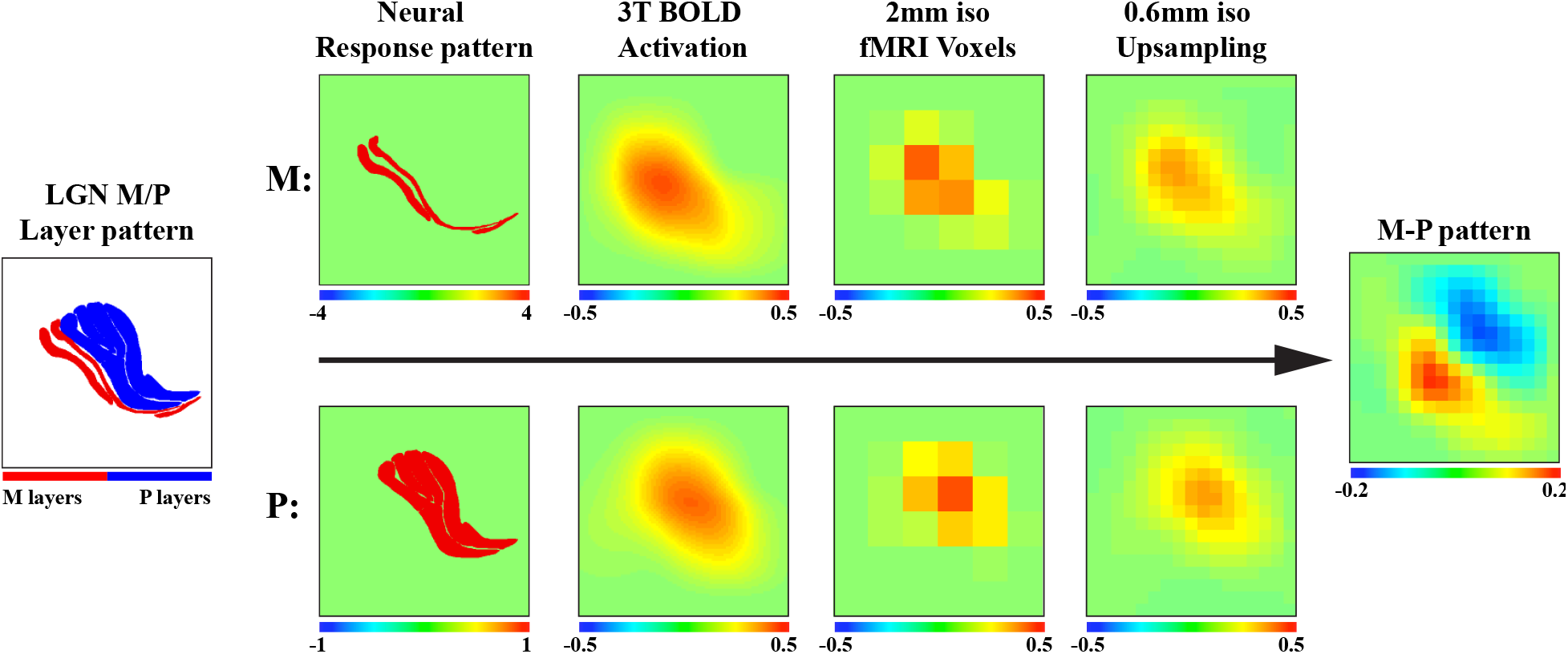
M-P pattern simulation for the 3T BOLD fMRI experiment.

**Figure S2.**
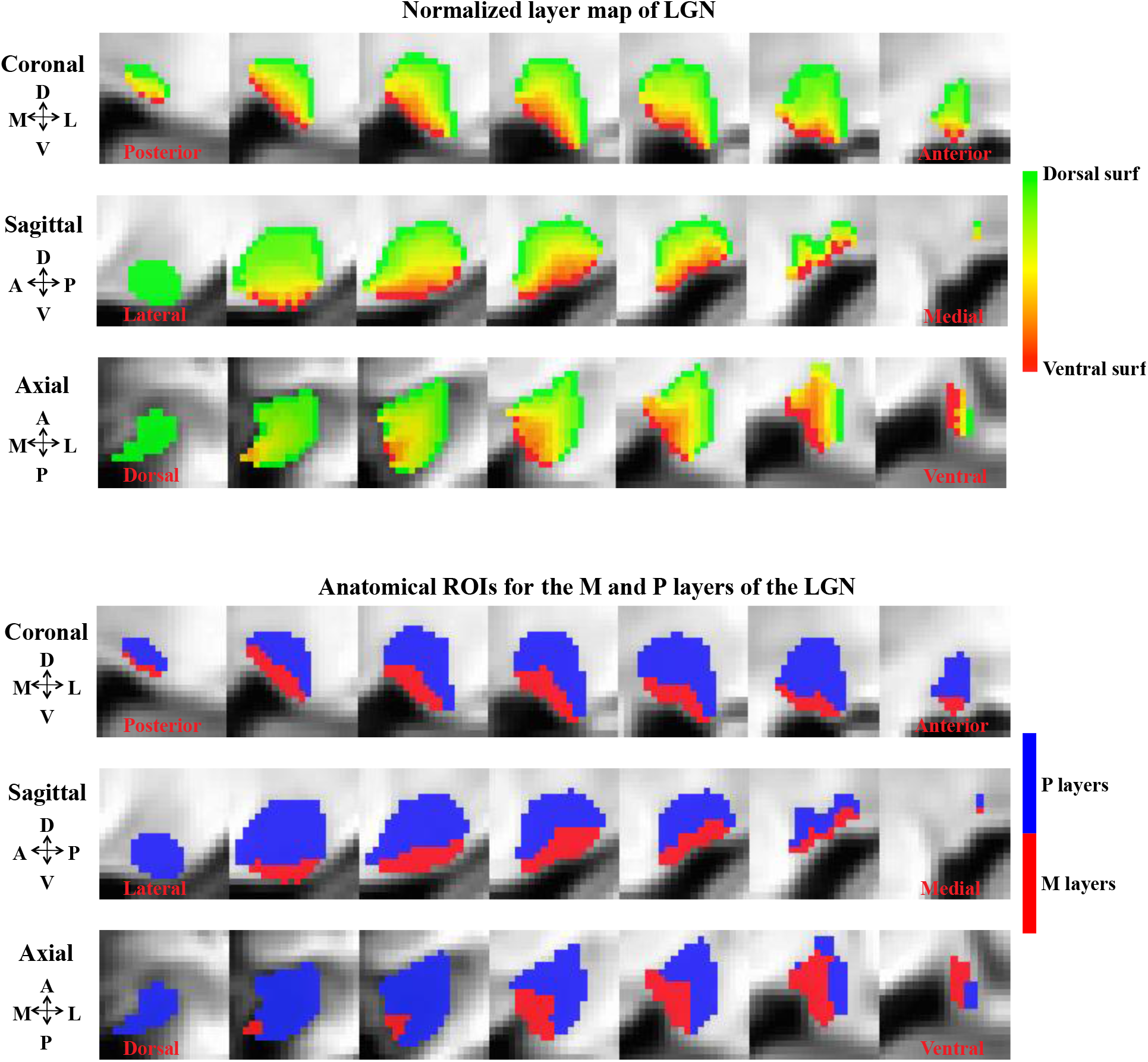
Normalized layer index map and ROIs for the M and P layers of the LGN (for the 7T experiment). Two layers of voxels were defined corresponding to the ventral and dorsal surfaces of the LGN. For the rest of voxels, we calculated a layer index as the normalized distances to the dorsal surface (ranged from 0 to 1). The ROIs for the M and P layers were determined from the layer index map according the volume ratio of M/P layers of human LGNs (1:4)

**Figure S3.**
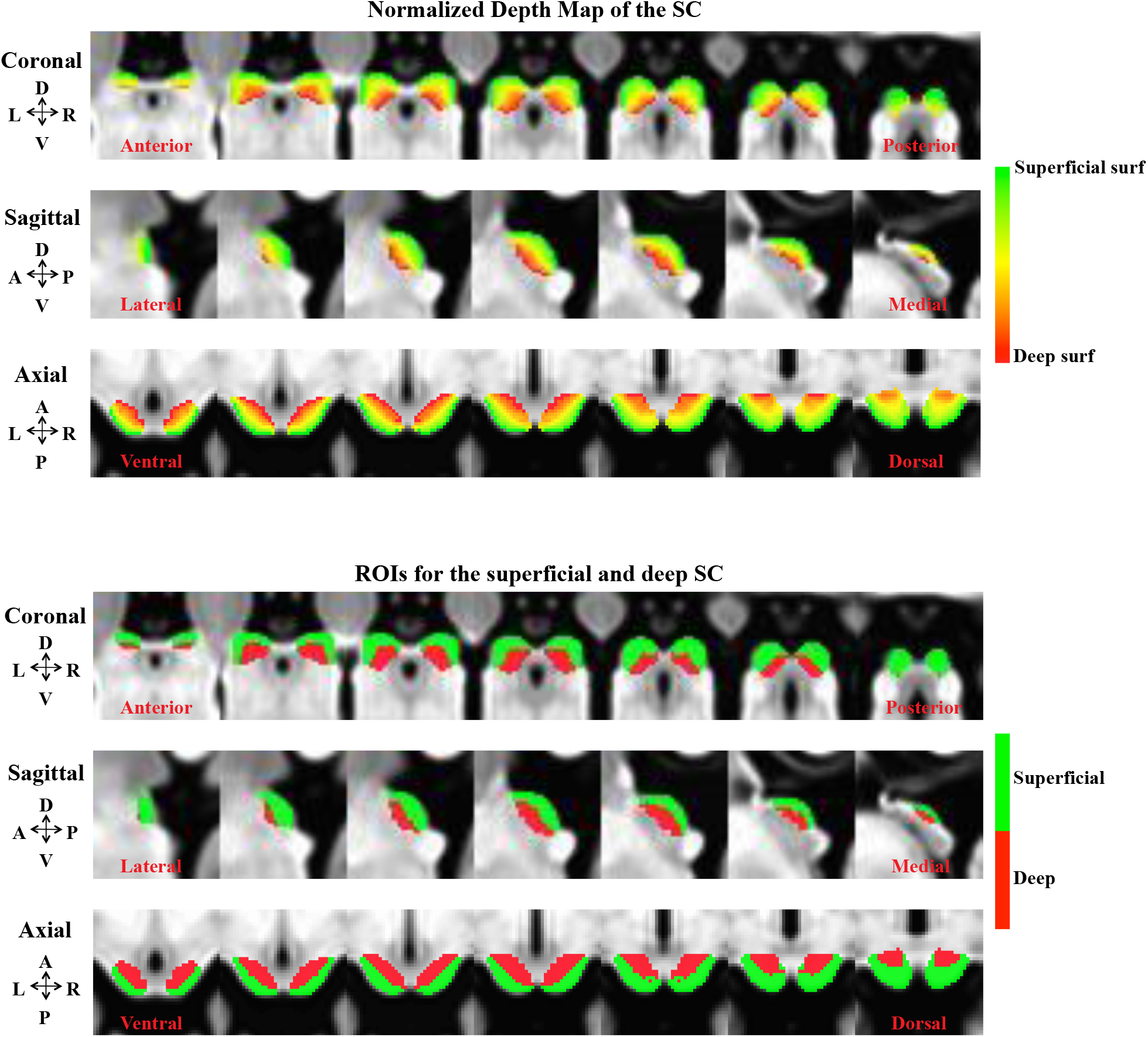
Normalized depth map and the superficial and deeper layers ROIs for the SC. Two layers of voxels were first defined from the superficial and deep surface of the SC. Then a normalized depth map was calculated for each voxel as the ratio of shortest distances to the superficial and deep surfaces (from 0/superficial to 1/deep). The volume of the SC was split into a superficial and a deeper part at depth = 0.5.

**Figure S4.**
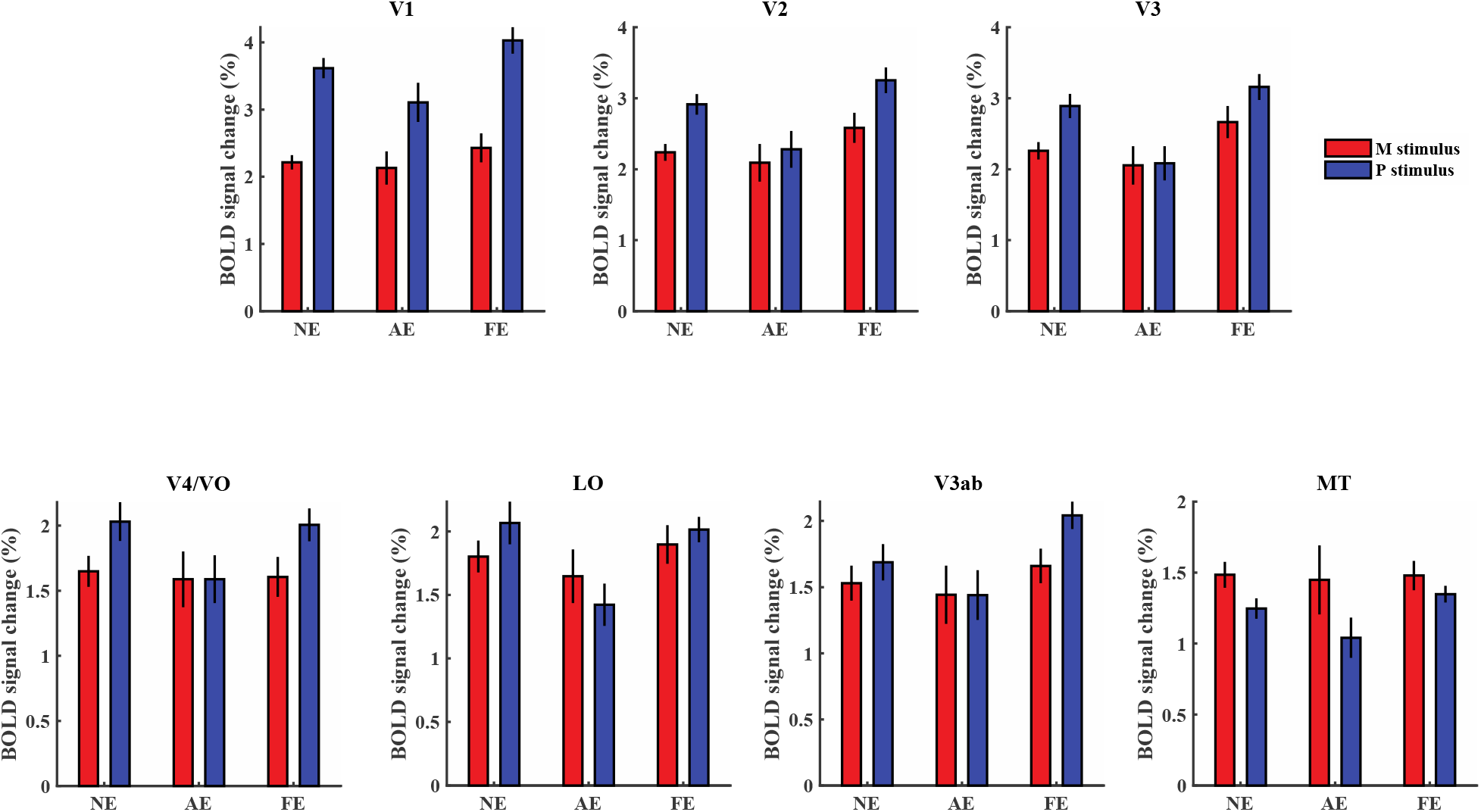
3T fMRI response to the M and P stimuli in cortical visual areas. Significant two-way interaction between eyes (AE/NE) and stimuli (M/P) was found in V1, V2, V3, V4/VO, and LO (all *p* < 0.05), but not in V3ab and MT (both *p* > 0.2).

**Figure S5.**
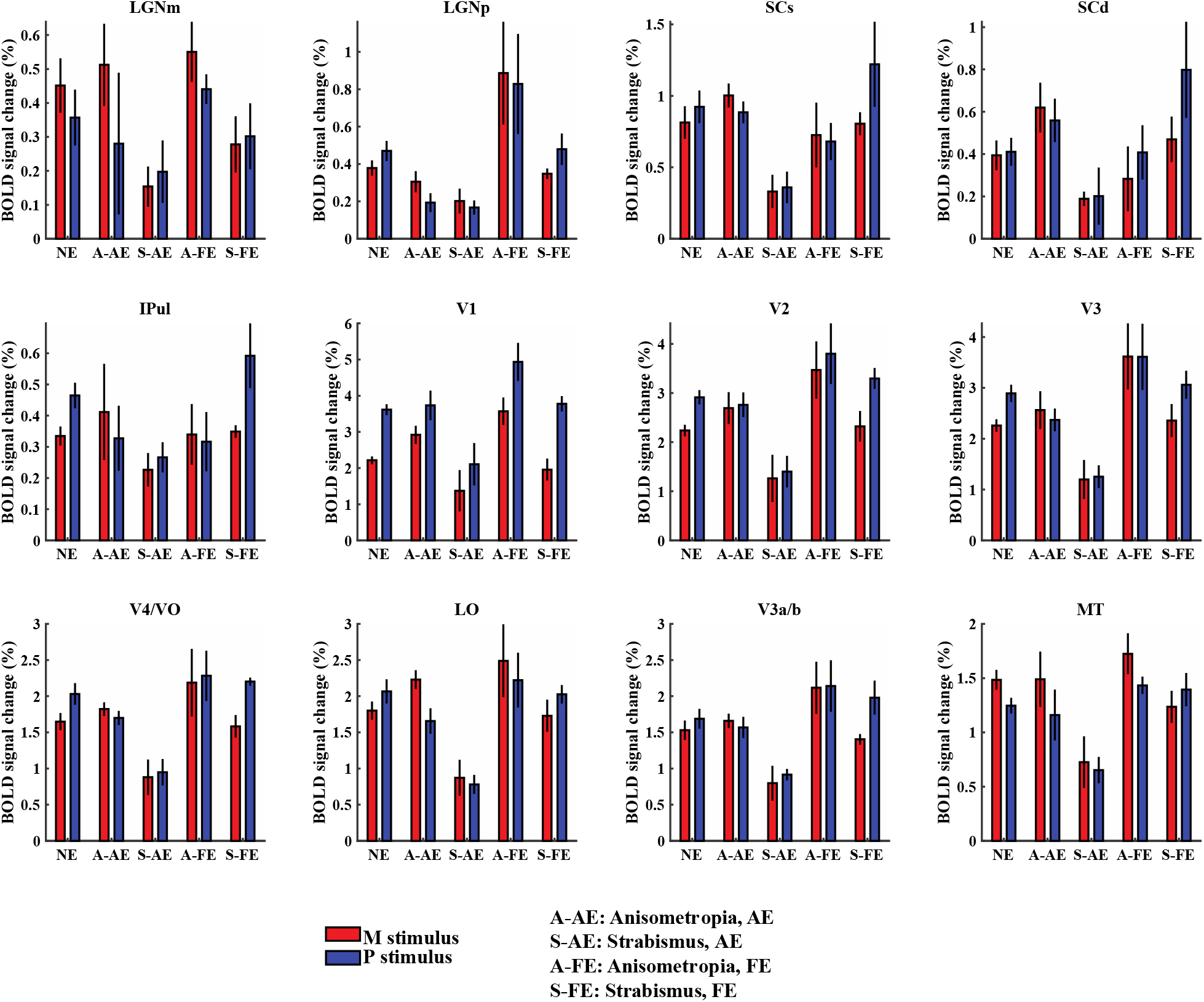
Subcortical and cortical responses for anisometropia and strabismus patients. Significant difference between the response to the amblyopic eye of Anisometropia and Strabismus patients can be found in higher order visual areas: V3, V4/VO, LO, and MT *(p* < 0.05 for interaction of groups (anisometropia/Strabismus) and eyes (AE/FE).

### Supplemental fMRI experiment: the influence of fixation instability on the BOLD response of early visual areas

**Figure S6.**
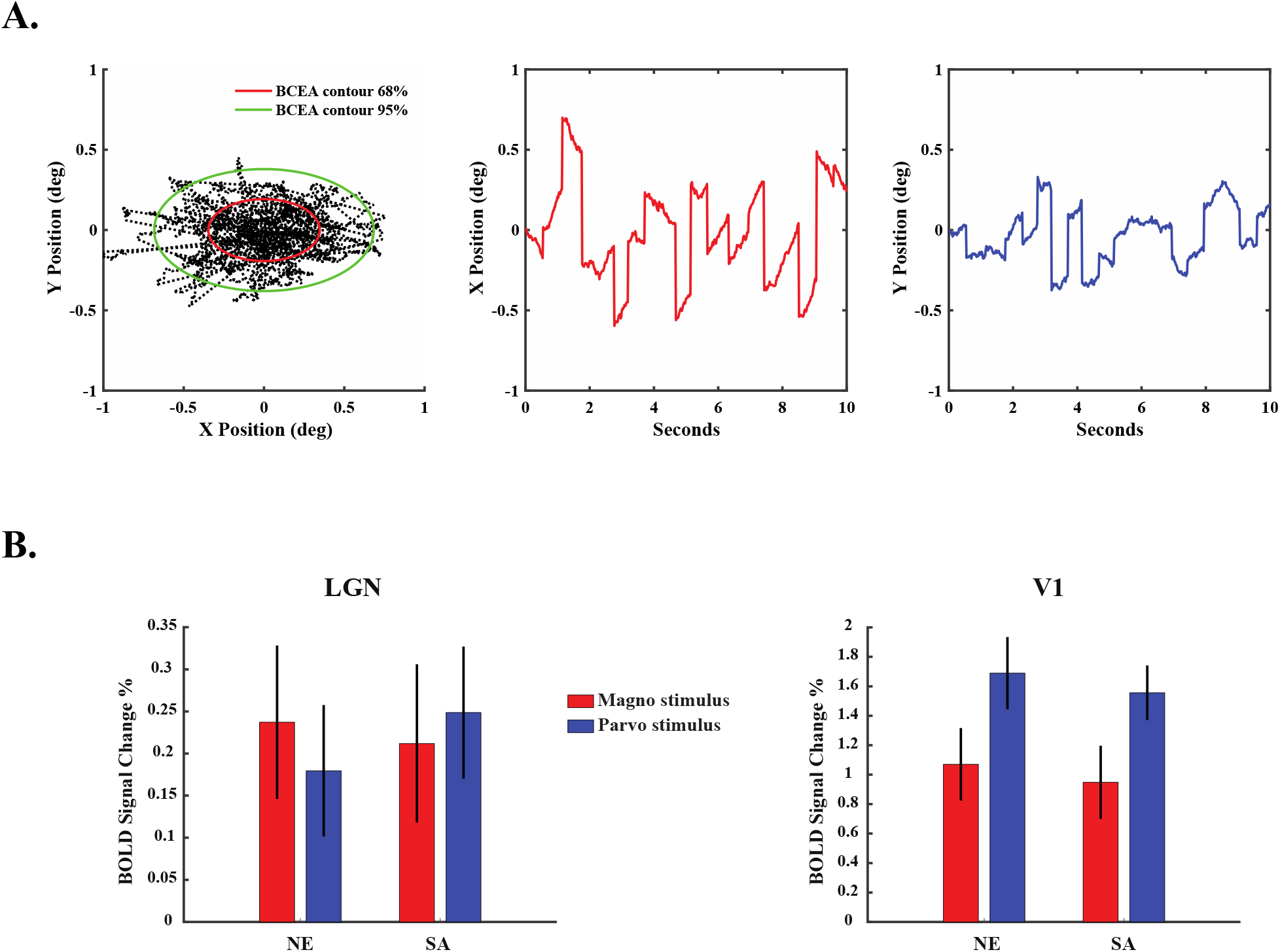
(A) Simulated fixational eye movements in the strabismus eye viewing condition. The left figure shows the simulated eye positions. The 68% and 95% BCEAs (bivariate contour ellipse area) are indicated by the red and green ellipse contours, respectively. The middle and the right figures represent the horizontal (X) and vertical (Y) eye movements. **(B) BOLD response of the LGN (left) and V1 (right) to M and P stimuli in the normal eye viewing (NE) and simulated strabismus eye viewing conditions (SA).**

It is known that amblyopia have abnormal fixational eye movements, especially those with strabismus. To investigate the influence of fixation instability on BOLD responses of early visual areas, we performed a control fMRI experiment on normal subjects. During the experiment, subjects viewed the M- and P-biased visual stimuli with and without jerky motion to simulate the retinal motion of unsteady fixational eye movements of strabismus. The non-dominate eye with correct refractive errors was tested, and the opposing eye was covered by a black eye patch. Examples of simulated eye movement traces were shown in Figure S6A. The simulation matched the characteristics of eye movements of strabismus from a previous study (Chung, Kumar et al. 2015): area of 68% bivariate contour ellipse (BCEA, 0.21 deg^2^), microsaccade frequency (2.9 Hz), microsaccade amplitude (26.9 arc min), slow drift speed (2.64 deg/sec), slow drift amplitude (11.2 arc min). Subjects completed three fMRI runs for the normal viewing condition (NE), and three runs for the strabismus eye viewing condition (SA). Each run consists of 160 functional volumes, with 2 seconds acquisition time for each volume. The subjective isoluminance of red and green was adjusted for each individual with a minimal flicker procedure, repeated four times in the scanner before the functional scans. Subjects were instructed to maintain fixation during the experiment. Eight healthy subjects with corrected to normal vision participated in the study (5 females, age 31.3±4.4 years).

MRI data were acquired with a 3T scanner (Siemens, Prisma) with a 20-channel phasearray coil (NOVA medical) from Beijing MRI center for Brain Research (BMCBR), at the Institute of Biophysics, Chinese Academy of Sciences. Functional images were acquired with a gradient echo planar imaging (GE-EPI) sequence (2mm isotropic voxels, 27 axial slices, 96×96 matrix with 2mm in-plane resolution, 2mm thickness, TR/TE = 2000/28 ms, flip angle = 90 deg). A T1-MPRAGE sequence was used to acquire anatomical images (1mm isotropic voxels, 224 sagittal 1mm-thick slices, 256 × 256 matrix with 1mm in-plane resolution, TR/TE = 2600/3.02 ms, inversion time = 900 ms, flip angle = 8°).

ROIs of the LGN were defined by an experienced researcher based on T1 contrast from T1-MPRAGE anatomical images, in which the LGN appeared darker than surrounding tissues. ROIs of V1 were defined on the cortical surface based on stimulus activation (M+P) and Benson 2018 HCP retinotopic atlas. BOLD responses to the M and P stimuli in the LGN and V1 were shown in Figure S6B. Two-way repeated-measures ANOVA with within-subject factors of stimuli (M/P stimuli) and eyes (normal/strabismus eyes) showed no significant main effect of eyes on the BOLD response of the LGN (F(1,7) = 0.35, *p* = 0.573) and V1(F(1,7) = 0.896, *p* = 0.375). Post-hoc t-test also showed no significant difference between the normal and strabismus eye viewing conditions for either M or P stimuli *(p* > 0.05 for all comparisons). These results suggest that reduced BOLD activity of the LGN was unlikely due to fixation instability of the amblyopic eye.

**Table S1.**
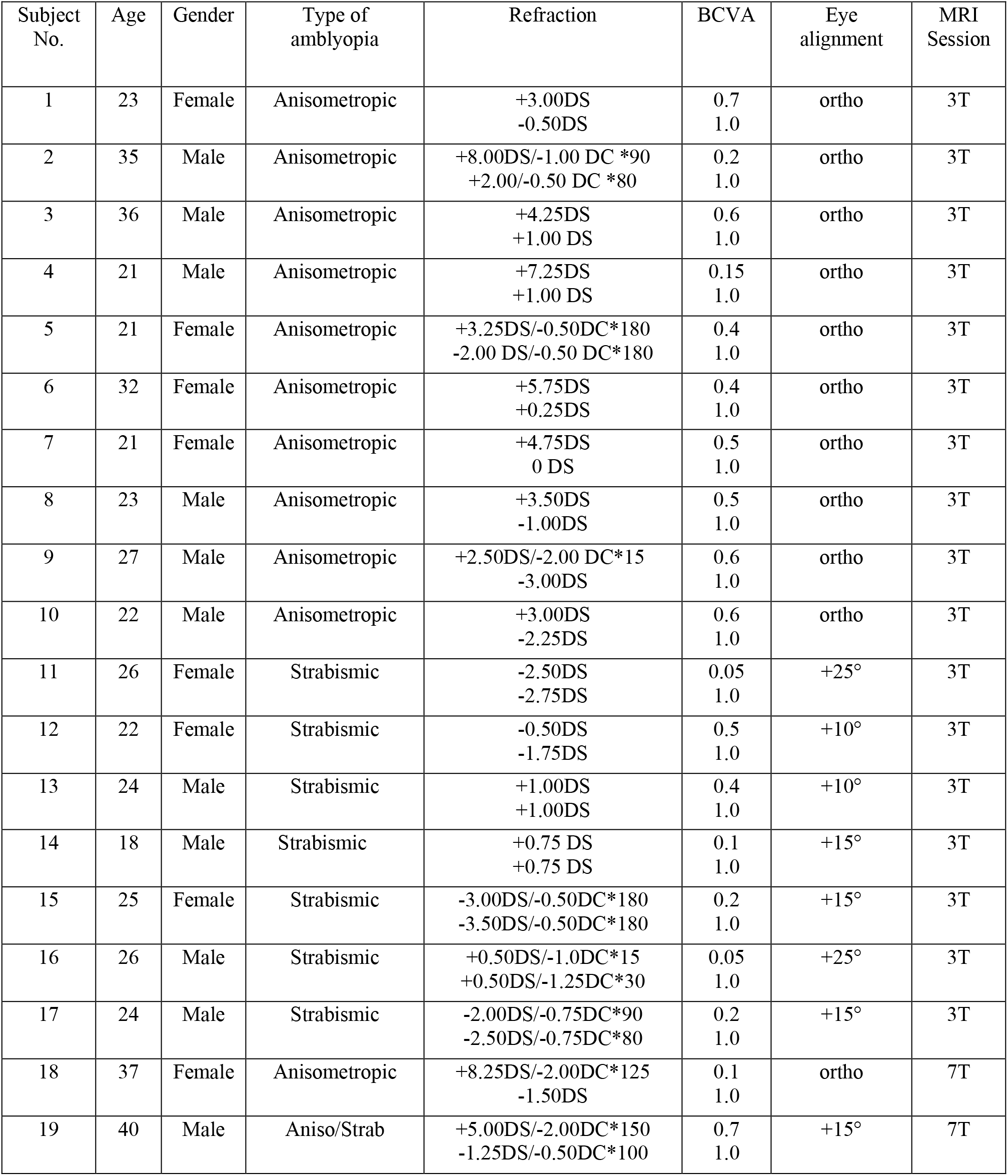
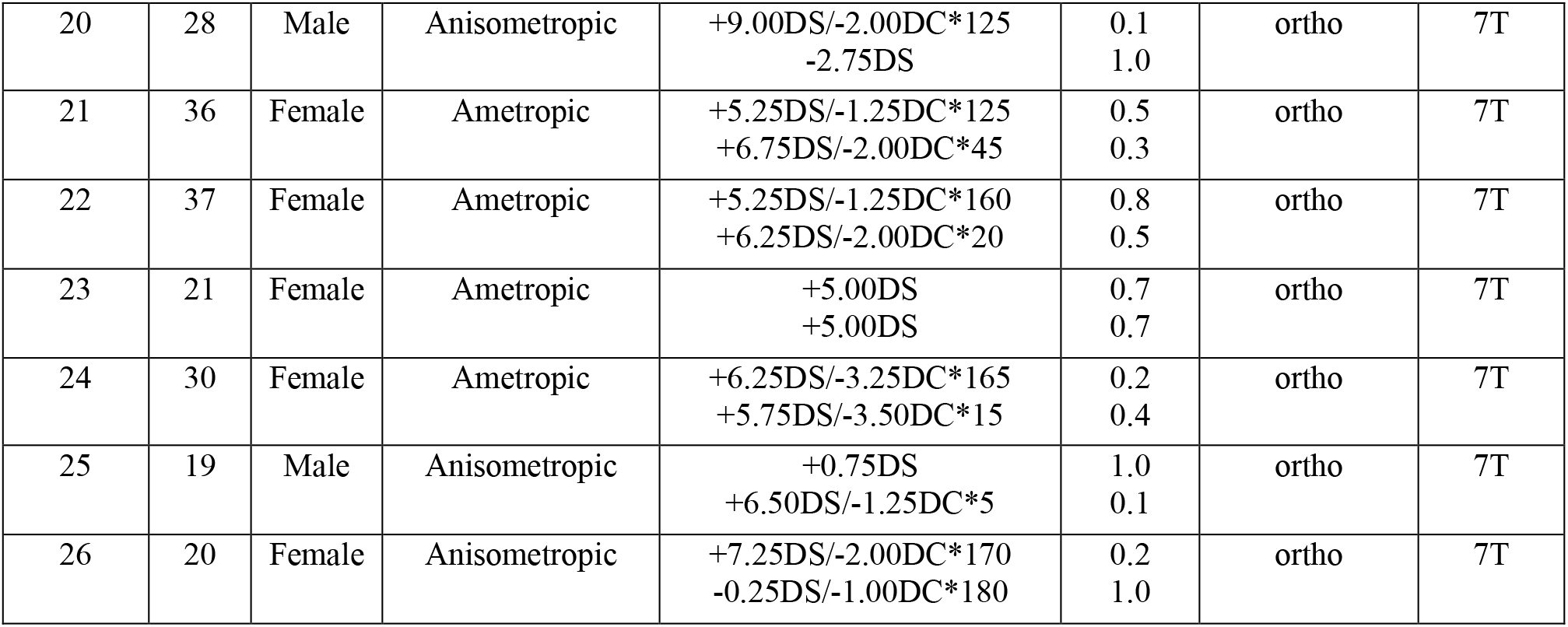
Clinical characteristics of amblyopia patients

| Subject No. | Age | Gender | Type of amblyopia | Refraction | BCVA | Eye alignment | MRI Session |
| --- | --- | --- | --- | --- | --- | --- | --- |
| 1 | 23 | Female | Anisometropic | +3.00DS<br>-0.50DS | 0.7<br>1.0 | ortho | 3T |
| 2 | 35 | Male | Anisometropic | +8.00DS/-1.00 DC *90<br>+2.00/-0.50 DC *80 | 0.2<br>1.0 | ortho | 3T |
| 3 | 36 | Male | Anisometropic | +4.25DS<br>+1.00 DS | 0.6<br>1.0 | ortho | 3T |
| 4 | 21 | Male | Anisometropic | +7.25DS<br>+1.00 DS | 0.15<br>1.0 | ortho | 3T |
| 5 | 21 | Female | Anisometropic | +3.25DS/-0.50DC*180<br>-2.00 DS/-0.50 DC*180 | 0.4<br>1.0 | ortho | 3T |
| 6 | 32 | Female | Anisometropic | +5.75DS<br>+0.25DS | 0.4<br>1.0 | ortho | 3T |
| 7 | 21 | Female | Anisometropic | +4.75DS<br>0 DS | 0.5<br>1.0 | ortho | 3T |
| 8 | 23 | Male | Anisometropic | +3.50DS<br>-1.00DS | 0.5<br>1.0 | ortho | 3T |
| 9 | 27 | Male | Anisometropic | +2.50DS/-2.00 DC*15<br>-3.00DS | 0.6<br>1.0 | ortho | 3T |
| 10 | 22 | Male | Anisometropic | +3.00DS<br>-2.25DS | 0.6<br>1.0 | ortho | 3T |
| 11 | 26 | Female | Strabismic | -2.50DS<br>-2.75DS | 0.05<br>1.0 | +25° | 3T |
| 12 | 22 | Female | Strabismic | -0.50DS<br>-1.75DS | 0.5<br>1.0 | +10° | 3T |
| 13 | 24 | Male | Strabismic | +1.00DS<br>+1.00DS | 0.4<br>1.0 | +10° | 3T |
| 14 | 18 | Male | Strabismic | +0.75 DS<br>+0.75 DS | 0.1<br>1.0 | +15° | 3T |
| 15 | 25 | Female | Strabismic | -3.00DS/-0.50DC*180<br>-3.50DS/-0.50DC*180 | 0.2<br>1.0 | +15° | 3T |
| 16 | 26 | Male | Strabismic | +0.50DS/-1.0DC*15<br>+0.50DS/-1.25DC*30 | 0.05<br>1.0 | +25° | 3T |
| 17 | 24 | Male | Strabismic | -2.00DS/-0.75DC*90<br>-2.50DS/-0.75DC*80 | 0.2<br>1.0 | +15° | 3T |
| 18 | 37 | Female | Anisometropic | +8.25DS/-2.00DC*125<br>-1.50DS | 0.1<br>1.0 | ortho | 7T |
| 19 | 40 | Male | Aniso/Strab | +5.00DS/-2.00DC*150<br>-1.25DS/-0.50DC*100 | 0.7<br>1.0 | +15° | 7T |
| 20 | 28 | Male | Anisometropic | +9.00DS/-2.00DC*125<br>-2.75DS | 0.1<br>1.0 | ortho | 7T |
| 21 | 36 | Female | Ametropic | +5.25DS/-1.25DC*125<br>+6.75DS/-2.00DC*45 | 0.5<br>0.3 | ortho | 7T |
| 22 | 37 | Female | Ametropic | +5.25DS/-1.25DC*160<br>+6.25DS/-2.00DC*20 | 0.8<br>0.5 | ortho | 7T |
| 23 | 21 | Female | Ametropic | +5.00DS<br>+5.00DS | 0.7<br>0.7 | ortho | 7T |
| 24 | 30 | Female | Ametropic | +6.25DS/-3.25DC*165<br>+5.75DS/-3.50DC*15 | 0.2<br>0.4 | ortho | 7T |
| 25 | 19 | Male | Anisometropic | +0.75DS<br>+6.50DS/-1.25DC*5 | 1.0<br>0.1 | ortho | 7T |
| 26 | 20 | Female | Anisometropic | +7.25DS/-2.00DC*170<br>-0.25DS/-1.00DC*180 | 0.2<br>1.0 | ortho | 7T |

## References

Abbie, A. A. (1933). The Blood Supply of the Lateral Geniculate Body, with a Note on the Morphology of the Choroidal Arteries. J Anat, 67(Pt 4), 491–521. https://www.ncbi.nlm.nih.gov/pubmed/17104443

Adams, M. M., Hof, P. R., Gattass, R., Webster, M. J., & Ungerleider, L. G. (2000). Visual cortical projections and chemoarchitecture of macaque monkey pulvinar. J Comp Neurol, 419(3), 377–393. https://doi.org/10.1002/(sici)1096-9861(20000410)419:3<377::aid-cne9>3.0.co;2-e

Allen, B., Spiegel, D. P., Thompson, B., Pestilli, F., & Rokers, B. (2015). Altered white matter in early visual pathways of humans with amblyopia. Vision Research, 114, 48–55. https://doi.org/10.1016/j.visres.2014.12.021

Amunts, K., Lepage, C., Borgeat, L., Mohlberg, H., Dickscheid, T., Rousseau, M. E., Bludau, S., Bazin, P. L., Lewis, L. B., Oros-Peusquens, A. M., Shah, N. J., Lippert, T., Zilles, K., & Evans, A. C. (2013). BigBrain: an ultrahigh-resolution 3D human brain model. Science, 340(6139), 1472–1475. https://doi.org/10.1126/science.1235381

Anderson, S. J., & Swettenham, J. B. (2006). Neuroimaging in human amblyopia. Strabismus, 14(1), 21–35. https://doi.org/10.1080/09273970500538082

Andrews, T. J., Halpern, S. D., & Purves, D. (1997). Correlated size variations in human visual cortex, lateral geniculate nucleus, and optic tract. Journal of Neuroscience, 17(8), 2859–2868. http://www.ncbi.nlm.nih.gov/pubmed/9092607

Arcaro, M. J., Pinsk, M. A., & Kastner, S. (2015). The Anatomical and Functional Organization of the Human Visual Pulvinar. Journal of Neuroscience, 35(27), 9848–9871. https://doi.org/10.1523/JNEUROSCI.1575-14.2015

Avants, B. B., Tustison, N. J., Song, G., Cook, P. A., Klein, A., & Gee, J. C. (2011). A reproducible evaluation of ANTs similarity metric performance in brain image registration. Neuroimage, 54(3), 2033–2044. https://doi.org/10.1016/j.neuroimage.2010.09.025

Barnes, G. R., Hess, R. F., Dumoulin, S. O., Achtman, R. L., & Pike, G. B. (2001). The cortical deficit in humans with strabismic amblyopia. J Physiol, 555(Pt 1), 281–297. https://doi.org/10.1111/j.1469-7793.2001.0281b.x

Beckstead, R. M., & Frankfurter, A. (1982). The distribution and some morphological features of substantia nigra neurons that project to the thalamus, superior colliculus and pedunculopontine nucleus in the monkey. Neuroscience, 7(10), 2377–2388. https://doi.org/10.1016/0306-4522(82)90202-0

Benson, N. C., Jamison, K. W., Arcaro, M. J., Vu, A. T., Glasser, M. F., Coalson, T. S., Van Essen, D. C., Yacoub, E., Ugurbil, K., Winawer, J., & Kay, K. (2018). The Human Connectome Project 7 Tesla retinotopy dataset: Description and population receptive field analysis. J Vis, 18(13), 23. https://doi.org/10.1167/18.13.23

Berman, R. A., & Wurtz, R. H. (2010). Functional identification of a pulvinar path from superior colliculus to cortical area MT. Journal of Neuroscience, 50(18), 6342–6354. https://doi.org/10.1523/JNEUROSCI.6176-09.2010

Brainard, D. H. (1997). The Psychophysics Toolbox. Spat Vis, 10(4), 433–436. http://www.ncbi.nlm.nih.gov/entrez/query.fcgi?cmd=Retrieve&db=PubMed&dopt=Citation&list_uids=9176952

Briggs, F., & Usrey, W. M. (2011). Corticogeniculate feedback and visual processing in the primate. J Physiol, 589(Pt 1), 33–40. https://doi.org/10.1113/jphysiol.2010.193599

Choi, M. Y., Lee, K. M., Hwang, J. M., Choi, D. G., Lee, D. S., Park, K. H., & Yu, Y. S. (2001). Comparison between anisometropic and strabismic amblyopia using functional magnetic resonance imaging. British Journal of Ophthalmology, 85(9), 1052–1056. https://doi.org/DOI10.1136/bjo.85.9.1052

Chung, S. T., Kumar, G., Li, R. W., & Levi, D. M. (2015). Characteristics of fixational eye movements in amblyopia: Limitations on fixation stability and acuity? Vision Res, 114, 87–99. https://doi.org/10.1016/j.visres.2015.01.016

Connolly, M., & Van Essen, D. (1984). The representation of the visual field in parvicellular and magnocellular layers of the lateral geniculate nucleus in the macaque monkey. J Comp Neurol, 226(4), 544–564. https://doi.org/10.1002/cne.902260408

Cox, R. W. (1996). AFNI: software for analysis and visualization of functional magnetic resonance neuroimages. Comput Biomed Res, 29(3), 162–173. http://www.ncbi.nlm.nih.gov/entrez/query.fcgi?cmd=Retrieve&db=PubMed&dopt=Citation&list_uids=8812068

Derrington, A. M., Krauskopf, J., & Lennie, P. (1984). Chromatic mechanisms in lateral geniculate nucleus of macaque. J Physiol, 357, 241–265. https://www.ncbi.nlm.nih.gov/pubmed/6512691

Derrington, A. M., & Lennie, P. (1984). Spatial and temporal contrast sensitivities of neurones in lateral geniculate nucleus of macaque. J Physiol, 357, 219–240. http://www.ncbi.nlm.nih.gov/entrez/query.fcgi?cmd=Retrieve&db=PubMed&dopt=Citation&list_uids=6512690

DeSimone, K., Viviano, J. D., & Schneider, K. A. (2015). Population Receptive Field Estimation Reveals New Retinotopic Maps in Human Subcortex. Journal of Neuroscience, 35(27), 9836–9847. https://doi.org/10.1523/JNEUROSCI.3840-14.2015

Felleman, D. J., & Van Essen, D. C. (1991). Distributed hierarchical processing in the primate cerebral cortex. Cereb Cortex, 1(1), 1–47. https://www.ncbi.nlm.nih.gov/pubmed/1822724

Finlay, B. L., Schiller, P. H., & Volman, S. F. (1976). Quantitative studies of single-cell properties in monkey striate cortex. IV. Corticotectal cells. J Neurophysiol, 39(6), 1352–1361. https://doi.org/10.1152/jn.1976.39.6.1352

Fries, W. (1984). Cortical projections to the superior colliculus in the macaque monkey: a retrograde study using horseradish peroxidase. J Comp Neurol, 230(1), 55–76. http://www.ncbi.nlm.nih.gov/entrez/query.fcgi?cmd=Retrieve&db=PubMed&dopt=Citation&list_uids=6096414

Friston, K. J., Harrison, L., & Penny, W. (2003). Dynamic causal modelling. Neuroimage, 19(4), 1273–1302. https://www.ncbi.nlm.nih.gov/pubmed/12948688

Friston, K. J., Litvak, V., Oswal, A., Razi, A., Stephan, K. E., van Wijk, B. C. M., Ziegler, G., & Zeidman, P. (2016). Bayesian model reduction and empirical Bayes for group (DCM) studies. Neuroimage, 128, 413–431. https://doi.org/10.1016/j.neuroimage.2015.11.015

Gati, J. S., Menon, R. S., Ugurbil, K., & Rutt, B. K. (1997). Experimental determination of the BOLD field strength dependence in vessels and tissue. Magn Reson Med, 38(2), 296–302. https://www.ncbi.nlm.nih.gov/pubmed/9256111

Gonzalez, E. G., Wong, A. M., Niechwiej-Szwedo, E., Tarita-Nistor, L., & Steinbach, M. J. (2012). Eye position stability in amblyopia and in normal binocular vision. Invest Ophthalmol Vis Sci, 53(9), 5386–5394. https://doi.org/10.1167/iovs.12-9941

Heidemann, R. M., Anwander, A., Feiweier, T., Knosche, T. R., & Turner, R. (2012). k-space and q-space: Combining ultra-high spatial and angular resolution in diffusion imaging using ZOOPPA at 7 T. Neuroimage, 60(2), 967–978. https://doi.org/10.1016/j.neuroimage.2011.12.081

Hess, R. F., & Baker Jr., C. L. (1984). Assessment of retinal function in severely amblyopic individuals. Vision Res, 24(10), 1367–1376. https://doi.org/10.1016/0042-6989(84)90192-5

Hess, R. F., Li, X. F., Mansouri, B., Thompson, B., & Hansen, B. C. (2009). Selectivity as well as Sensitivity Loss Characterizes the Cortical Spatial Frequency Deficit in Amblyopia. Human Brain Mapping, 50(12), 4054–4069. https://doi.org/10.1002/hbm.20829

Hess, R. F., Li, X., Lu, G., Thompson, B., & Hansen, B. C. (2010). The contrast dependence of the cortical fMRI deficit in amblyopia; a selective loss at higher contrasts. Hum Brain Mapp, 51(8), 1233–1248. https://doi.org/10.1002/hbm.20931

Hess, R. F., Thompson, B., Gole, G. A., & Mullen, K. T. (2010). The amblyopic deficit and its relationship to geniculo-cortical processing streams. J Neurophysiol, 104(1), 475–483. https://doi.org/10.1152/jn.01060.2009

Hess, R. F., Thompson, B., Gole, G., & Mullen, K. T. (2009). Deficient responses from the lateral geniculate nucleus in humans with amblyopia. Eur J Neurosci, 29(5), 1064–1070. https://doi.org/10.1111/j.1460-9568.2009.06650.x

Hubel, D. H., & Livingstone, M. S. (1990). Color and contrast sensitivity in the lateral geniculate body and primary visual cortex of the macaque monkey. Journal of Neuroscience, 10(7), 2223–2237. https://www.ncbi.nlm.nih.gov/pubmed/2198331

Hubel, D. H., Wiesel, T. N., & LeVay, S. (1977). Plasticity of ocular dominance columns in monkey striate cortex. Philos Trans R Soc LondB Biol Sci, 278(961), 377–409. https://doi.org/10.1098/rstb.1977.0050

Jay, M. F., & Sparks, D. L. (1987). Sensorimotor integration in the primate superior colliculus. II. Coordinates of auditory signals. J Neurophysiol, 57(1), 35–55. https://doi.org/10.1152/jn.1987.57.1.35

Joly, O., & Franko, E. (2014). Neuroimaging of amblyopia and binocular vision: a review. Front Integr Neurosci, 8, 62. https://doi.org/10.3389/fnint.2014.00062

Kim, S. G. (2018). Biophysics of BOLD fMRI investigated with animal models. J Magn Reson. https://doi.org/10.1016/j.jmr.2018.04.006

Kriegeskorte, N., Simmons, W. K., Bellgowan, P. S., & Baker, C. I. (2009). Circular analysis in systems neuroscience: the dangers of double dipping. Nat Neurosci, 12(5), 535–540. https://doi.org/10.1038/nn.2303

Kuang, X., Poletti, M., Victor, J. D., & Rucci, M. (2012). Temporal encoding of spatial information during active visual fixation. Curr Biol, 22(6), 510–514. https://doi.org/10.1016/j.cub.2012.01.050

Lerner, Y., Hendler, T., Malach, R., Harel, M., Leiba, H., Stolovitch, C., & Pianka, P. (2006). Selective fovea-related deprived activation in retinotopic and high-order visual cortex of human amblyopes. Neuroimage, 55(1), 169–179. https://doi.org/10.1016/j.neuroimage.2006.06.026

Levi, D. M. (2006). Visual processing in amblyopia: human studies. Strabismus, 14(1), 11–19. https://doi.org/10.1080/09273970500536243

Levi, D. M., & Harwerth, R. S. (1977). Spatio-temporal interactions in anisometropic and strabismic amblyopia. Invest Ophthalmol Vis Sci, 16(1), 90–95. https://www.ncbi.nlm.nih.gov/pubmed/832970

Li, H., Yang, X., Gong, Q., Chen, H., Liao, M., & Liu, L. (2013). BOLD responses to different temporospatial frequency stimuli in V1 and V2 visual cortex of anisometropic amblyopia. Eur J Ophthalmol, 23(2), 147–155. https://doi.org/10.5301/ejo.5000211

Lock, T. M., Baizer, J. S., & Bender, D. B. (2003). Distribution of corticotectal cells in macaque. Exp Brain Res, 151(4), 455–470. https://doi.org/10.1007/s00221-003-1500-y

Lyon, D. C., Nassi, J. J., & Callaway, E. M. (2010). A disynaptic relay from superior colliculus to dorsal stream visual cortex in macaque monkey. Neuron, 65(2), 270–279. https://doi.org/10.1016/j.neuron.2010.01.003

Mai, J. K., Majtanik, M., & Paxinos, G. (2015). Atlas of the human brain. Academic Press.

May, P. J. (2006). The mammalian superior colliculus: laminar structure and connections. Neuroanatomy of the Oculomotor System, 151, 321–378. https://doi.org/10.1016/S0079-6123(05)51011-2

Movshon, J. A., Eggers, H. M., Gizzi, M. S., Hendrickson, A. E., Kiorpes, L., & Boothe, R. G. (1987). Effects of early unilateral blur on the macaque’s visual system. III. Physiological observations. Journal of Neuroscience, 7(5), 1340–1351. https://www.ncbi.nlm.nih.gov/pubmed/3572484

Pauli, W. M., Nili, A. N., & Tyszka, J. M. (2018). A high-resolution probabilistic in vivo atlas of human subcortical brain nuclei. Sci Data, 5, 180063. https://doi.org/10.1038/sdata.2018.63

Pelli, D. G. (1997). The VideoToolbox software for visual psychophysics: transforming numbers into movies. Spat Vis, 10(4), 437–442. http://www.ncbi.nlm.nih.gov/entrez/query.fcgi?cmd=Retrieve&db=PubMed&dopt=Citation&list_uids=9176953

Perry, V. H., & Cowey, A. (1984). Retinal ganglion cells that project to the superior colliculus and pretectum in the macaque monkey. Neuroscience, 12(4), 1125–1137. https://doi.org/10.1016/0306-4522(84)90007-1

Purushothaman, G., Marion, R., Li, K., & Casagrande, V. A. (2012). Gating and control of primary visual cortex by pulvinar. Nat Neurosci, 15(6), 905–912. https://doi.org/10.1038/nn.3106

Qi, S., Mu, Y. F., Cui, L. B., Li, R., Shi, M., Liu, Y., Xu, J. Q., Zhang, J., Yang, J., & Yin, H. (2016). Association of Optic Radiation Integrity with Cortical Thickness in Children with Anisometropic Amblyopia. Neuroscience Bulletin, 32(1), 51–60. https://doi.org/10.1007/s12264-015-0005-6

Qian, Y., Zou, J., Zhang, Z., An, J., Zuo, Z., Zhuo, Y., Wang, D. J. J., & Zhang, P. (2020). Robust functional mapping of layer-selective responses in human lateral geniculate nucleus with high-resolution 7T fMRI. Proceedings of the Royal Society B: Biological Sciences, 287(1925). https://doi.org/10.1098/rspb.2020.0245

Rucci, M., Iovin, R., Poletti, M., & Santini, F. (2007). Miniature eye movements enhance fine spatial detail. Nature, 447(7146), 851–854. https://doi.org/10.1038/nature05866

Saalmann, Y. B., Pinsk, M. A., Wang, L., Li, X., & Kastner, S. (2012). The Pulvinar Regulates Information Transmission Between Cortical Areas Based on Attention Demands. Science, 557(6095), 753–756. https://doi.org/10.1126/science.1223082

Schlesinger, B. (2012). The upper brainstem in the human: its nuclear configuration and vascular supply. Springer Science & Business Media.

Schneider, K. A., Richter, M. C., & Kastner, S. (2004). Retinotopic organization and functional subdivisions of the human lateral geniculate nucleus: a high-resolution functional magnetic resonance imaging study. Journal of Neuroscience, 24(41), 8975–8985. http://www.ncbi.nlm.nih.gov/entrez/query.fcgi?cmd=Retrieve&db=PubMed&dopt=Citation&list_uids=15483116

Shaikh, A. G., Otero-Millan, J., Kumar, P., & Ghasia, F. F. (2016). Abnormal Fixational Eye Movements in Amblyopia. PLoS One, 11(3), e0149953. https://doi.org/10.1371/journal.pone.0149953

Sherman, S. M., & Guillery, R. W. (2002). The role of the thalamus in the flow of information to the cortex. Philos Trans R Soc Lond B Biol Sci, 557(1428), 1695–1708. https://doi.org/10.1098/rstb.2002.1161

Shipp, S. (2003). The functional logic of cortico-pulvinar connections. Philos Trans R Soc Lond B Biol Sci, 558(1438), 1605–1624. https://doi.org/10.1098/rstb.2002.1213

Subramanian, V., Jost, R. M., & Birch, E. E. (2013). A quantitative study of fixation stability in amblyopia. Invest Ophthalmol Vis Sci, 54(3), 1998–2003. https://doi.org/10.1167/iovs.12-11054

Thompson, B., Villeneuve, M. Y., Casanova, C., & Hess, R. F. (2012). Abnormal cortical processing of pattern motion in amblyopia: evidence from fMRI. Neuroimage, 60(2), 1307–1315. https://doi.org/10.1016/j.neuroimage.2012.01.078

Ungerleider, L. G., Desimone, R., Galkin, T. W., & Mishkin, M. (1984). Subcortical projections of area MT in the macaque. J Comp Neurol, 225(3), 368–386. https://doi.org/10.1002/cne.902230304

Verghese, P., McKee, S. P., & Levi, D. M. (2019). Attention deficits in Amblyopia. Current Opinion in Psychology, 29, 199–204. https://doi.org/10.1016/j.copsyc.2019.03.011

Wang, W. F., Kiyosawa, M., Ishiwata, K., & Mochizuki, M. (2005). Glucose metabolism in the visual structures of rat monocularly deprived by eyelid suture after postnatal eye opening. Japanese Journal of Ophthalmology, 49(1), 6–11. https://doi.org/10.1007/s10384-004-0146-z

White, B. J., Berg, D. J., Kan, J. Y., Marino, R. A., Itti, L., & Munoz, D. P. (2017). Superior colliculus neurons encode a visual saliency map during free viewing of natural dynamic video. Nat Commun, 8, 14263. https://doi.org/10.1038/ncomms14263

White, B. J., Boehnke, S. E., Marino, R. A., Itti, L., & Munoz, D. P. (2009). Color-related signals in the primate superior colliculus. Journal of Neuroscience, 29(39), 12159–12166. https://doi.org/10.1523/JNEUROSCI.1986-09.2009

White, B. J., Kan, J. Y., Levy, R., Itti, L., & Munoz, D. P. (2017). Superior colliculus encodes visual saliency before the primary visual cortex. Proceedings of the National Academy of Sciences of the United States of America, 114(35), 9451–9456. https://doi.org/10.1073/pnas.1701003114

Wiberg, M., Westman, J., & Blomqvist, A. (1987). Somatosensory projection to the mesencephalon: an anatomical study in the monkey. J Comp Neurol, 264(1), 92–117. https://doi.org/10.1002/cne.902640108

Wiesel, T. N., & Hubel, D. H. (1963). Effects of Visual Deprivation on Morphology and Physiology of Cells in the Cats Lateral Geniculate Body. J Neurophysiol, 26, 978–993. https://doi.org/10.1152/jn.1963.26.6.978

Xie, S., Gong, G. L., Xiao, J. X. I., Ye, J. T., Liu, H. H., Gan, X. L., Jiang, Z. T., & Jiang, X. X. (2007). Underdevelopment of Optic Radiation in Children With Amblyopia: A Tractography Study. American Journal of Ophthalmology, 143(4), 642–646. https://doi.org/10.1016/j.ajo.2006.12.009

Yan, Y., Zhaoping, L., & Li, W. (2018). Bottom-up saliency and top-down learning in the primary visual cortex of monkeys. Proc Natl Acad Sci U SA, 115(41), 10499–10504. https://doi.org/10.1073/pnas.1803854115

Yu, Q., Zhang, P., Qiu, J., & Fang, F. (2016). Perceptual Learning of Contrast Detection in the Human Lateral Geniculate Nucleus. Current Biology, 26(23). https://doi.org/10.1016/j.cub.2016.09.034

Zeidman, P., Jafarian, A., Corbin, N., Seghier, M. L., Razi, A., Price, C. J., & Friston, K. J. (2019). A guide to group effective connectivity analysis, part 1: First level analysis with DCM for fMRI. Neuroimage, 200, 174–190. https://doi.org/10.1016/j.neuroimage.2019.06.031

Zeidman, P., Jafarian, A., Seghier, M. L., Litvak, V., Cagnan, H., Price, C. J., & Friston, K. J. (2019). A guide to group effective connectivity analysis, part 2: Second level analysis with PEB. Neuroimage, 200, 12–25. https://doi.org/10.1016/j.neuroimage.2019.06.032

Zele, A. J., Pokorny, J., Lee, D. Y., & Ireland, D. (2007). Anisometropic amblyopia: spatial contrast sensitivity deficits in inferred magnocellular and parvocellular vision. Invest Ophthalmol Vis Sci, 48(8), 3622–3631. https://doi.org/10.1167/iovs.06-1207

Zele, Andrew J., Wood, J. M., & Girgenti, C. C. (2010). Magnocellular and parvocellular pathway mediated luminance contrast discrimination in amblyopia. Vision Res, 50(10), 969–976. https://doi.org/10.1016/j.visres.2010.03.002

Zhang, P., Wen, W., Sun, X., & He, S. (2015). Selective reduction of fMRI responses to transient achromatic stimuli in the magnocellular layers of the LGN and the superficial layer of the SC of early glaucoma patients. Hum Brain Mapp, 57(2). https://doi.org/10.1002/hbm.23049

Zhang, P., Zhou, H., Wen, W., & He, S. (2015). Layer-specific response properties of the human lateral geniculate nucleus and superior colliculus. Neuroimage, 111. https://doi.org/10.1016/j.neuroimage.2015.02.025

Zhaoping, L. (2008). Attention capture by eye of origin singletons even without awareness—A hallmark of a bottom-up saliencymap in the primary visual cortex. J Vis, 8(5), 1–18. http://www.journalofvision.org/content/8/5/1.full.pdf

Zhou, H., Schafer, R. J., & Desimone, R. (2016). Pulvinar-Cortex Interactions in Vision and Attention. Neuron, 89(1), 209–220. https://doi.org/10.1016/j.neuron.2015.11.034

